# HLApollo: A superior transformer model for pan-allelic peptide-MHC-I presentation prediction, with diverse negative coverage, deconvolution and protein language features

**DOI:** 10.1101/2022.12.08.519673

**Authors:** William John Thrift, Nicolas W. Lounsbury, Quade Broadwell, Amy Heidersbach, Emily Freund, Yassan Abdolazimi, Qui T Phung, Jieming Chen, Aude-Hélène Capietto, Ann-Jay Tong, Christopher M. Rose, Craig Blanchette, Jennie R Lill, Benjamin Haley, Lélia Delamarre, Richard Bourgon, Kai Liu, Suchit Jhunjhunwala

## Abstract

Antigen presentation on MHC class I (MHC-I) is key to the adaptive immune response to cancerous cells. Computational prediction of peptide presentation by MHC-I has enabled individualized cancer immunotherapies. Here, we introduce HLApollo, a transformer-based approach with end-to-end modeling of MHC-I sequence, deconvolution, and flanking sequences. To achieve this, we develop a novel training strategy, negative set switching, which greatly reduces overfitting to falsely presumed negatives that are necessarily found in presentation datasets. HLApollo shows a meaningful improvement compared to recent MHC-I models on peptide presentation (20.19% average precision (AP)) and immunogenicity (4.1% AP). As expected, adding gene expression boosts the performance of HLApollo. More interestingly, we show that introduction of features from a protein language model, ESM 1b, remarkably recoups much of the benefits of gene expression in absence of true expression measurements. Finally, we demonstrate excellent pan-allelic generalization, and introduce a framework for estimating the expected accuracy of HLApollo for untrained alleles. This guides the use of HLApollo in a clinical setting, where rare alleles may be observed in some subjects, particularly for underrepresented minorities.

## Introduction

Antigen presentation by MHC class I molecules (MHC-I) is crucial to alert the adaptive immune system to the presence of non-self moieties. Recognition of somatically mutated peptides (neoantigens) by CD8+ T cells on the tumor cell surface drives anti-tumor immunity^1–5^. Recently, individualized neoantigen specific therapies (iNeST) have been developed to target neoantigens presented at the tumor surface^6–9^. This approach relies on accurate prediction of peptide presentation by MHC-I for identification of patient-specific personalized neoantigens.

The antigen presentation pathway and machinery for MHC-I consists of several components, including peptide processing by the proteasome, and other cytosolic and endoplasmic reticulum (ER)-resident proteases, transport of peptides to ER through specialized complexes, and peptide loading onto MHC-I aided by chaperones^10–13^. MHC-I presented peptides (ligands), typically 8-14 amino acids long, can be very diverse in their sequence because of the polymorphic nature of the MHC locus, with thousands of possible MHC-I alleles being present in the population. Besides, the locus is also polygenic, with 3 genes encoding the alpha chain in humans (HLA-A, HLA-B and HLA-C). Each of an individual’s 3-6 MHC-I allotypes can bind and present up to 10,000 unique peptides forming a distinct ligand repertoire^14–17^.

Given the high diversity of MHC allotypes and the entire ligand repertoire (ligandome), it has been a long-standing interest in the scientific community to understand and computationally model the ligandome. One of the earliest approaches to do so employed position specific weight matrices to identify peptide binding motifs^18^. In the third decade since then, several advanced and performant approaches have been developed, including the neural network approach implemented in NetMHC’s suite of tools^19–21^. An explosion of ligandome data obtained using liquid chromatography–tandem mass spectrometry has allowed development of several advanced deep learning based methods (Supplementary Table 1). However, several heuristic choices continue to be made in these methods to deal with data complexity. For example, some methods are a mix of allele-specific models instead of a pan-allele generalized model that can predict for untrained alleles. Multi-allelic (MA) data, wherein there is no experimentally assigned association between a specific allotype and ligand, need to be deconvolved. Some methods only use single allele (SA) data from engineered mono-allelic cell lines, while other methods conduct an upfront heuristic deconvolution of MA data that is not modeled. Given the large amount of training data and the capability of deep learning approaches, especially of attention-based approaches^22–24^, we aimed to develop a more generic and performant approach, avoiding heuristics.

In this work we present HLApollo, a pan-allelic, transformer-based model for predicting peptide presentation by MHC-I. The model performs end-to-end modeling of MA deconvolution, peptide processing, and pan-allelic training. We introduce negative set switching, a training strategy to mitigate the impact of falsely presumed negatives, which are necessarily introduced during negative set simulation. Our baseline HLApollo, which uses only amino acid sequences of the peptide, flanks and MHC-I, achieves the best performance on presentation (20.19% average precision (AP) over NetMHCPan4.1-EL^25^), CD8 T cell response (4.1% AP over NetMHCPan4.1-EL, Wells et al. dataset^26^), and study holdout presentation datasets (7.9% more peptides recovered than SHERPA-EL^27^). When gene expression information is available, an extended HLAPollo + expression model achieves an AP of 23.93% greater on a negative peptide-augmented test set than the next best comparable approach, HLAthena^28^. We develop a new strategy of adding gene-level presentation relevant features from protein language models, which improves performance on CD8 T cell response (0.5% AP over baseline HLApollo, Schmidt et al dataset^29^). Finally, we introduce a linear regression model for predicting out-of-training allele performance using only a priori information and no training data. This predicts an increase of HLApollo genotype coverage on several underrepresented ethnicities, raising the number of covered alleles by 791, and can guide future ligandome collection to be more equitable.

## Results

### Development of a comprehensive Immune Peptidomics database

Though several public ligandome databases of peptide-MHC-I complexes (pMHC-I) exist, we needed to develop a novel schema and database that adequately reflects the workflow of a typical immune peptidomics experiment^30–33^. Thus, we developed mhcDB, a SQLite database of ligandome and paired bulk RNAseq data that enables quality control at the granularity of sample runs, harmonization of different data modalities, and meta-analyses of the original experiments. The schema provides strict tracking of metadata (e.g. study, donor/sample, HLA genotypes, analyte, treatment, fragmentation technique, search parameters, etc.) and biophysical data (e.g. peptide sequence, standardized peptide modifications, protein mapping, etc.) (Fig. la)(Supplementary Fig. 1).

**Figure 1.**
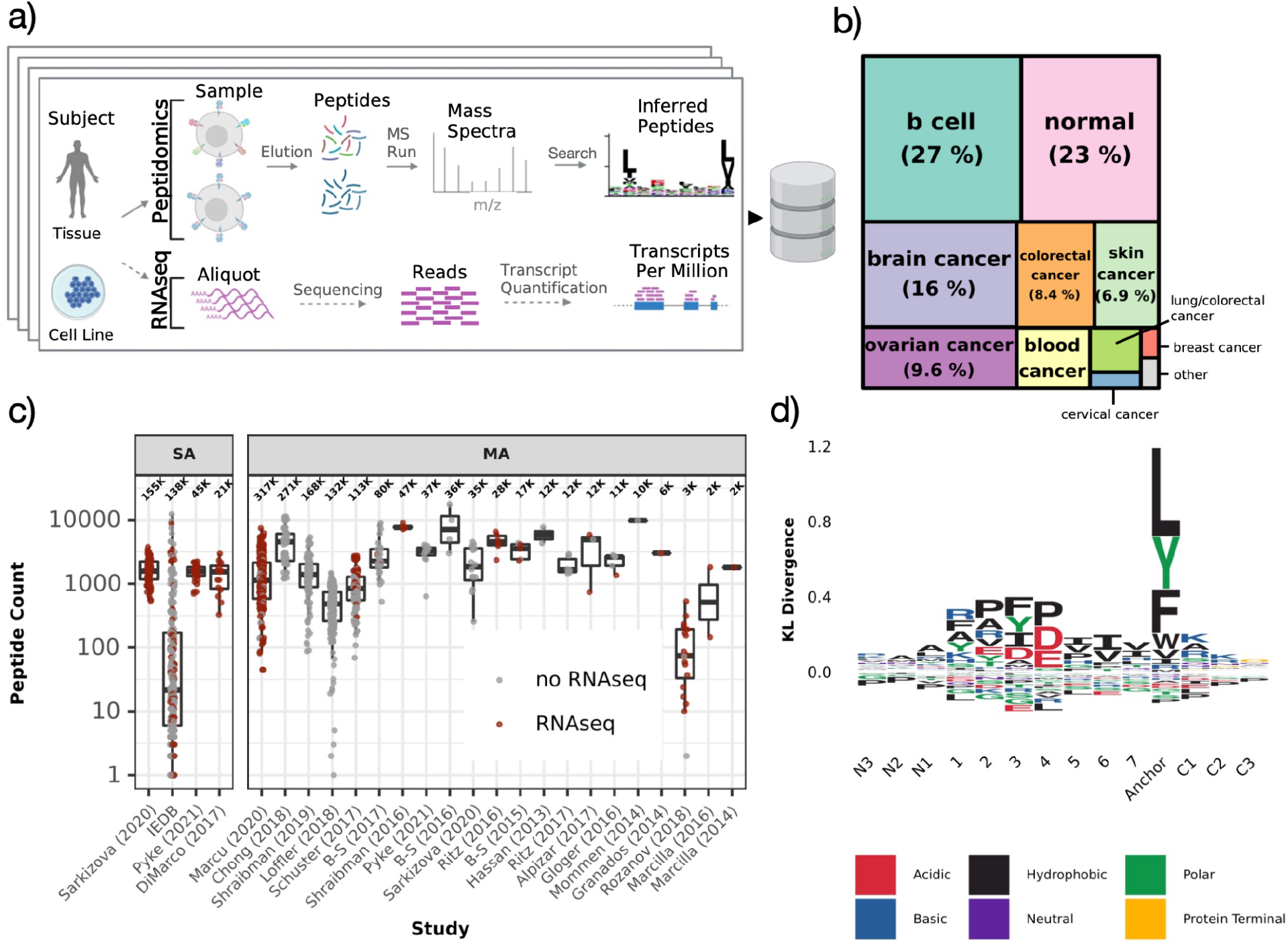
Summary of immune peptidomics data. a) Schematic of immunopeptidomics workflow and/or matched bulk RNAseq workflows for a given study. Data from 22 studies and IEDB was processed and imported into a sqlite database, mhcDB. b) Treemap plot showing the proportion of tissue types or disease indications across mhcDB. c) Unique peptide counts per sample from studies of single-allele (left) and multi-allele (right) data type, respectively. Samples with matched RNAseq are colored red. d) Position-wise amino acid enrichment at the N-terminal flank, peptide (positions 1-7, last position), C-terminal flank for peptides between 8 and 14 AA long, normalized by the number of peptides per HLA genotype.

We assembled a total of 953,693 unique {peptide, genotype} tuples across 347 unique HLA-I genotypes (SA + MA at 4-digit resolution), 305,646 unique peptides and 171 unique HLA-I alleles from a set of 22 published studies and IEDB^34^ (Fig. 1c)(Supplementary Table 2, 4)(see Supplementary methods). Data were filtered to retain WT peptides containing only canonical amino acids (no post-translational modifications) and a non-redundant set of studies in our database (Supplementary Fig. 2). Our ligandome data covers 8 cancer indications, while B cell lines, healthy tissue, brain cancer, colorectal cancer and skin cancer together account for ~80% of the data (Fig. 1b). We also collected available bulk RNAseq data from matched samples, subjects or healthy tissue, amounting to 43.8% of our ligandome data (Fig. 1c). We provide Supplementary Table 3 to track the datasets used to train and evaluate various models; the flagship benchmark (BM) dataset described above is referred to as BM1.

BM1’s (SA only) average positional probability matrix reveals patterns in line with previous studies^14,27,28^. In particular we note enrichment of hydrophobic/polar residues at the terminal anchor and acidic residues at auxiliary anchor positions 2-4 (Fig. 1d). Interestingly, the A/R/K ‘cleavage signal’ is enriched at C1 and P1.

In order to characterize the surface ligandome of our samples we plotted the frequency of top presenting genes along with properties such as expression, average protein length, and the distribution of presented peptides (Supplementary Fig. 3). Unsurprisingly, most of these genes are consistently highly expressed and/or encode long proteins, contributing to their high degree of presentation. Notably, 3 of the top 4 genes (*VIM, COL6A3, DYNC1H1*) consistently present across several indications and are necessary for the structural integrity of the cell. Moreover, the top presenting gene, *VIM*, is crucial for epithelial-to-mesenchymal transition, where its overexpression has been linked to metastasis in multiple cancer indications and poor prognosis in Acute Myeloid Leukemia^35–36^.

### HLApollo outperforms state of the art pMHC-I prediction methods

Fig. 2a depicts a schematic representation of our pan-allelic pMHC-I transformer^37^ architecture, HLApollo. HLApollo separately processes the peptide and MHC-I pseudo sequences before both representations are concatenated for further processing, ensuring good individual representations before relevant pMHC-I interactions are processed. HLApollo achieved an AP of 74.83% (74.67%-74.99% within one standard deviation) through 5-fold cross validation (see methods) on our test set BM1 (Supplementary Table 3).

**Figure 2.**
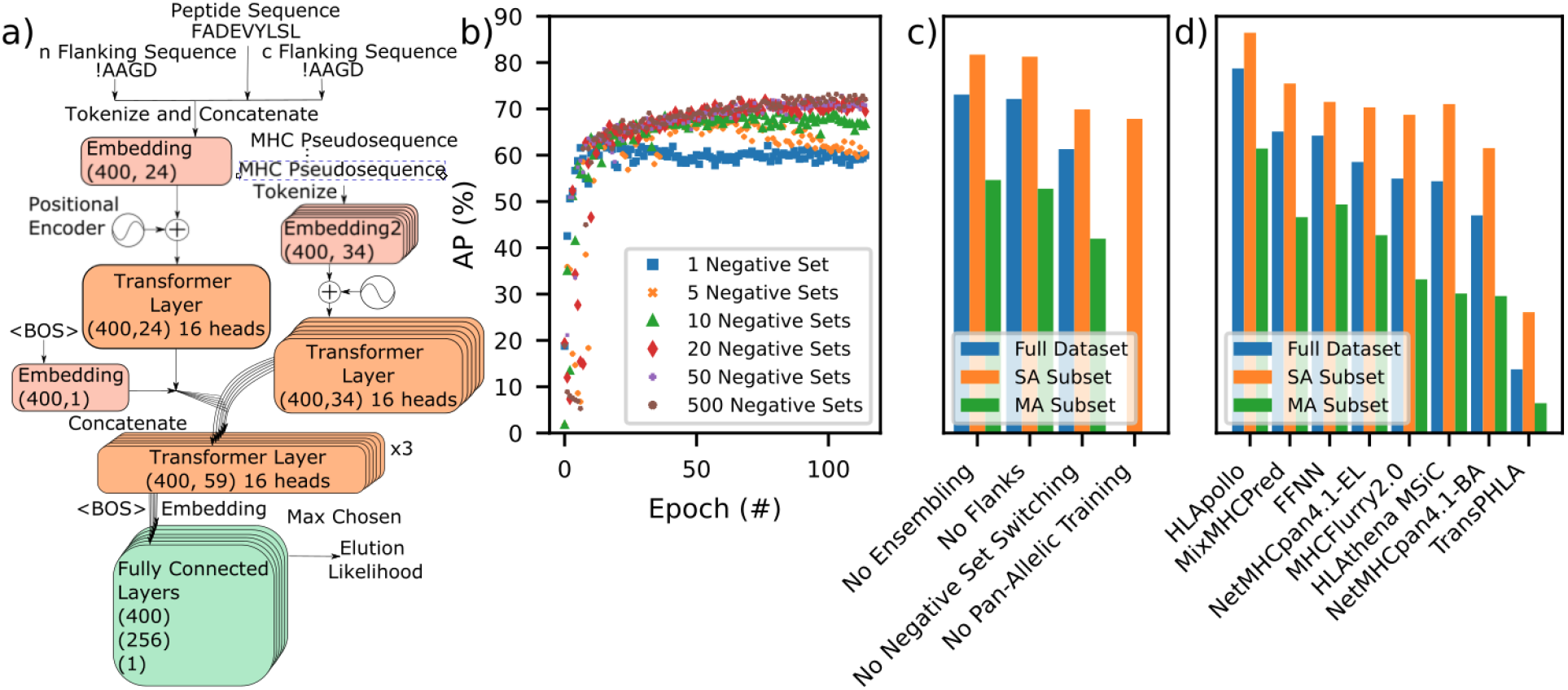
HLApollo model architecture and performance on dataset BM1. a) A schematic of the transformer-based model used throughout this work. BOS:Beginning of sequence b) Plot of the average precision over epochs for one, five, ten, twenty, fifty, and five hundred negative sets (blue square, orange x, green triangle, red diamond, purple cross, and brown circle, respectively) c) AP values of non-ensembled HLApollo and various individual ablations (all ablations performed without ensembling or 5 fold cross validation). Blue bars depict the performance on the full test set (MA and SA data), while orange and green bars depict the performance subset to just the SA and MA data, respectively. d) AP values of various contemporary models on the test dataset, and our implementation of a feed forward neural network (FFNN) model.

An important challenge of pMHC-I models is (over)fitting to falsely presumed negative peptides, which can occur in the training data due to sampling the reference proteome, but may actually be presented. A key innovation of this work is a new training strategy – negative set switching – where we sample a new exclusive set of negatives after each epoch of training, thereby preventing the model from observing false negatives more than once. The impact of the number of negative sets on performance is depicted in Fig. 2b. Overfitting was observed with ten or fewer negative sets. In all, the five hundred set negative set switching used here led to an improvement of 11.79% AP compared to one negative set, among the largest contributors in performance in the ablations we investigated.

Fig. 2c depicts the performance of HLApollo (without ensembling) and various ablations on BM1. Recently, HLAthena reported that a pan-allelic approach actually performed worse than individual models for their feed forward neural network (FFNN) model, especially for 8-mers. However, HLApollo was benefited by pan-allelic training across all peptide lengths, increasing AP by 13.9% (Fig. 2c, Supplementary Fig. 4). HLApollo jointly learns peptide processing, which we achieve by concatenating the peptide and flanking sequences (up to 5 residues) prior to being fed into the model. We further employ a regularization strategy: random removal of some or all of the flanking residues, to ensure good predictions on flanked, partially flanked, and flankless cases (see methods). End to end treatment of peptide processing leads to a 0.9% increase in AP.

Multiallelic (MA) data consists of more than twice as many peptides as SA data in our dataset, so it is important for models to learn effectively from such data. Unlike other approaches^27,38,39^, our model continuously learns which MHC-I allele presents a peptide by sending all MHC-I pseudo sequences through the model in parallel (Fig. 2a), and use the allele with maximum likelihood to evaluate loss. We note here that the performance gap between SA and MA data is smaller for HLApollo than other models considered in Fig. 2d.

For comparisons with other contemporary models, depicted in Fig. 2d), we ensembled 10 models for our flagship HLApollo, which led to an AP to 78.7%, 13.63% larger than any other model considered. In order to compare the use of a sequence based model with FFNNs common in the literature^25,28,40^, we trained our own, using a similar strategy as NetMHCpan. An FFNN architecture (with flank information, negative set switching, and ensembling) as opposed to an attentive sequence based architecture led to a large decline in AP of 14.5% (Fig. 2d).

### Inclusion of gene expression reduces false discovery at low expression levels

Besides peptide biophysical properties, expression of the source gene has been reported to be an important factor in determining presentation^28^. Empirical peptide enrichment (EPE) showed a log-linear relationship with expression (Fig. 3a). Interestingly, we observed EPE and HLApollo logit scores crossing 0 simultaneously. We also expect that expression especially reduces false positive calls in lowly or non-expressed genes. We switched to a 4999:1 negative:positive ratio test set to explore this effect (BM2). Due to the heterogeneity of expression values between samples we use sample-specific test sets to explore the impact of this effect. A logistic regression approach using HLApollo and expression increased AP by 6.03% (Fig. 3d).

**Figure 3.**
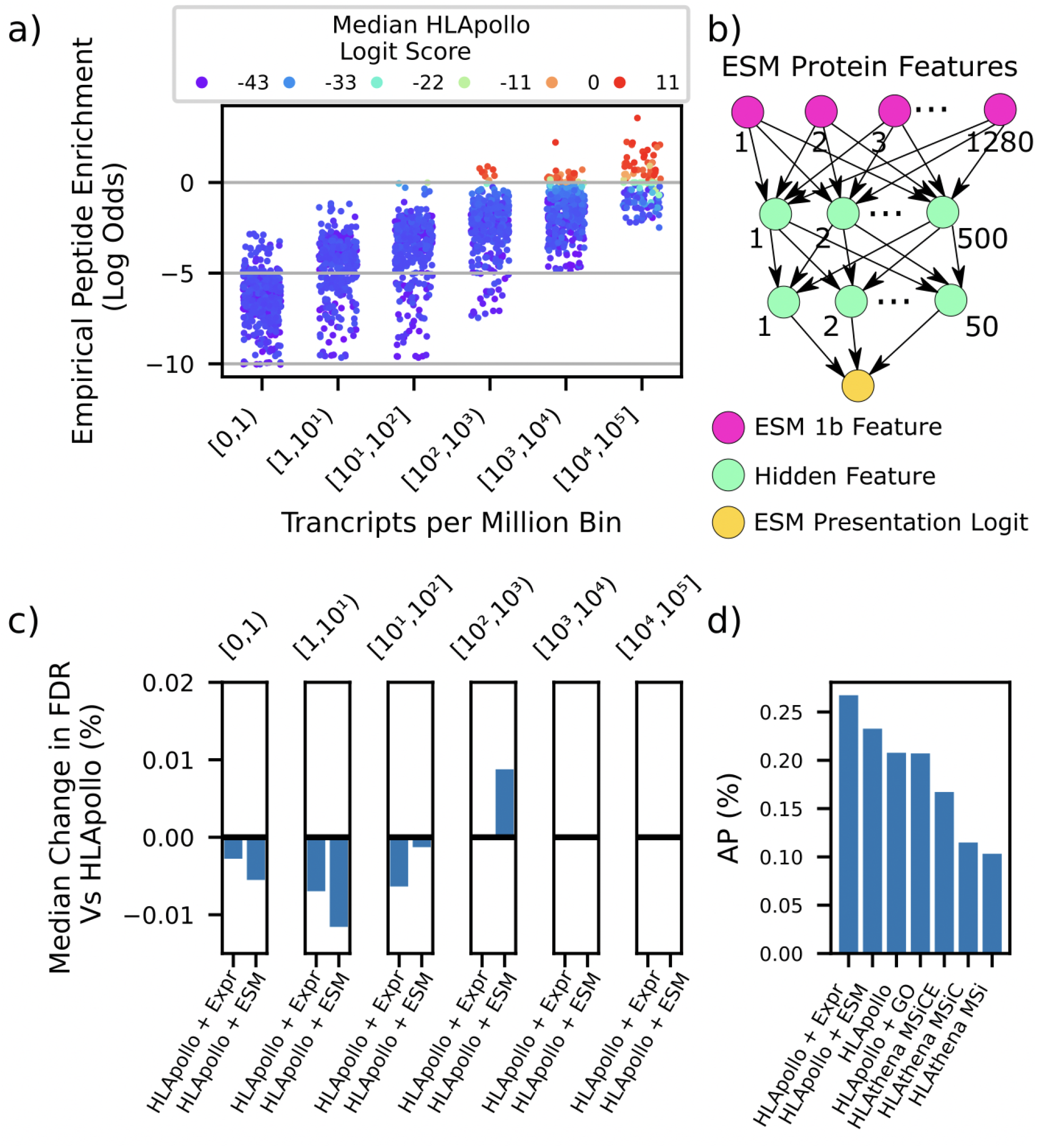
Gene expression or protein language model features boost HLApollo performance a) Binned expression (TPM) vs. empirical peptide enrichment (see Methods.) where each point represents a sample with a total of 50,000 peptides (combined positives and negatives). Samples (n=217) are colored by the median HLApollo logit prediction in a given expression bin. b) FFNN model used to extract presentation features from ESM1b protein features. c) median per-sample false discovery rate (FDR) gain over HLApollo of the logistic regression models further adding expression (left bars) or ESM_MHC-I_ score(right bars), at increasing expression levels. d) depicts the median performance of various extensions of HLApollo that utilize features beyond the sequence features considered in Fig. 2. All extensions were based on simple logistic regression using the baseline HLApollo score and the additional feature.

In addition to expression, previous studies showed that gene-specific features, such as protein localization annotation, such as that from Gene Ontology (GO), and presentation bias scores provide marginal improvement to presentation prediction^14,27,28^. Here, we sought to capture a wide range of gene-specific features at once by using protein language models. We used Evolutionary Scale Model (ESM) 1b^41^ to extract a gene-level embedding, from which an FFNN (depicted in Fig. 3b)) was trained to predict the presentation likelihood (ESM_MHC-I_). Using ESM_MHC-I_ with HLApollo in a logistic regression improved AP, while adding GO to HLApollo did not. However, we found that the ESM_MHC-I_ did not capture information about proteins that is orthogonal to protein expression: a logistic regression trained with HLApollo, ESM_MHC-I_, and expression actually performs 3.47% worse than HLApollo and expression, about the same as just HLApollo and ESM_MHC-I_ (data not shown in Fig. 3d)), and yielded a similar impact on FDR at different expression bins as HLApollo and expression (Fig 3c), suggesting that the gene-specific features captured by ESM_MHC-I_ are proxies for consistently lowly expressed genes. Regardless, these results show that ESM_MHC-I_ can be useful in applications where expression values are unknown, such as for important immunogenicity datasets considered below.

### Validation on tissue presentation holdouts and CD8-T cell response to neoantigens

Model generalization to unseen human tissue samples is paramount for clinical applications and challenged by the variety of biological/technical variation introduced therein. Fig. 4a–c compares HLApollo’s generalization to holdout human tissue studies (BM3) from Schuster et al.^42^, Löffler et al.^43^, and Pyke et al.^27^, respectively, with recent pMHC-I presentation models. Here we compared models’ study-holdout performance by the fraction of peptides that are ranked better than 0.1% percentile rank (see methods), as in other works^27,28^. HLApollo showed higher sensitivity than the other models considered in study holdout generalization.

**Figure 4.**
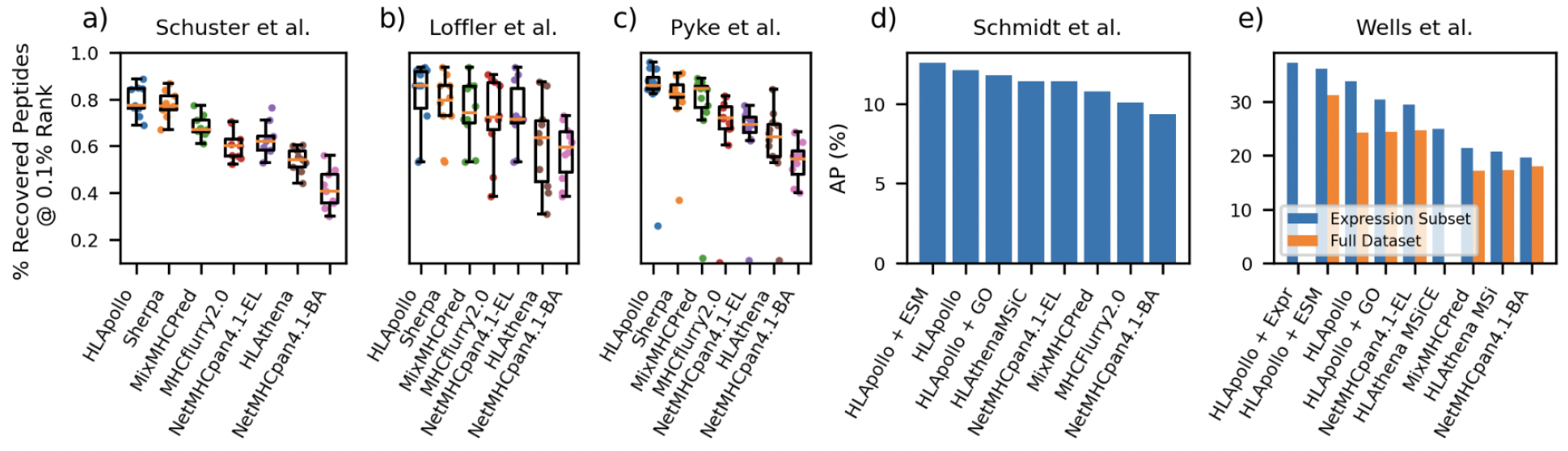
HLApollo shows superior performance in tissue holdout and immunogenicity test data a) depicts per sample boxplots of the fraction of eluted peptides ranked at less than 0.1% rank by various models for held out Schuster et al. b) Loffler et al., and c) Pyke et al. d) depicts the average AP of various models on data sets of pre-existing CD8 T cell response to cancer neoantigens, acquired from Schmidt et al. and e) average AP of various models on Wells et al after 500 bootstraps.

An important application of pMHC-I presentation models is to rank neoantigens for use in individualized neoantigen specific cancer therapies. To this end we considered cancer neoantigen immunogenicity datasets acquired / collected by Schmidt et al^44^ (BM4) and Wells et al^45^, (BM5) (Fig. 4d,e). HLApollo outperformed competing pMHC-I models for immunogenicity prediction. These immunogenicity datasets also demonstrate the utility of HLApollo + ESM-MHCI for predicting immunogenicity. We observe that HLApollo + ESM-MHCI improves AP by 0.5% and 2.0% for Schmidt et al. and Wells et al., respectively, compared to baseline HLApollo, while GO did not add any benefit.

### A bonafide deconvolution approach

Current deconvolution approaches for MA data require manual interpretations, or substantially rely on SA data-trained models to untangle MA data^27,38,46^. In contrast, our strategy enables *de novo* training from MA data. We rigorously evaluated our deconvolution strategy using a synthetic MA dataset (BM6), constructed from a ground-truth SA dataset. Borrowing from co-occurrence and exclusion principles^47^, we created ‘deconvolvable’ and ‘non-deconvolvable’ sets of genotypes (see methods). We compared a model trained on this synthetic MA dataset and one trained on the underlying SA data, evaluating both on SA test data. Remarkably, the model trained on MA data alone performed nearly identical to the model trained on SA data, with no distinction observed between ‘deconvolvable’, and ‘non-deconvolvable’ alleles (Fig. 5a). Overall, the model calls 93.9% of alleles correctly, based on the allele with the highest score from the MA-only model (91.1% for ‘non-deconvolvable’ and 95.6% for ‘deconvolvable’).

**Figure 5.**
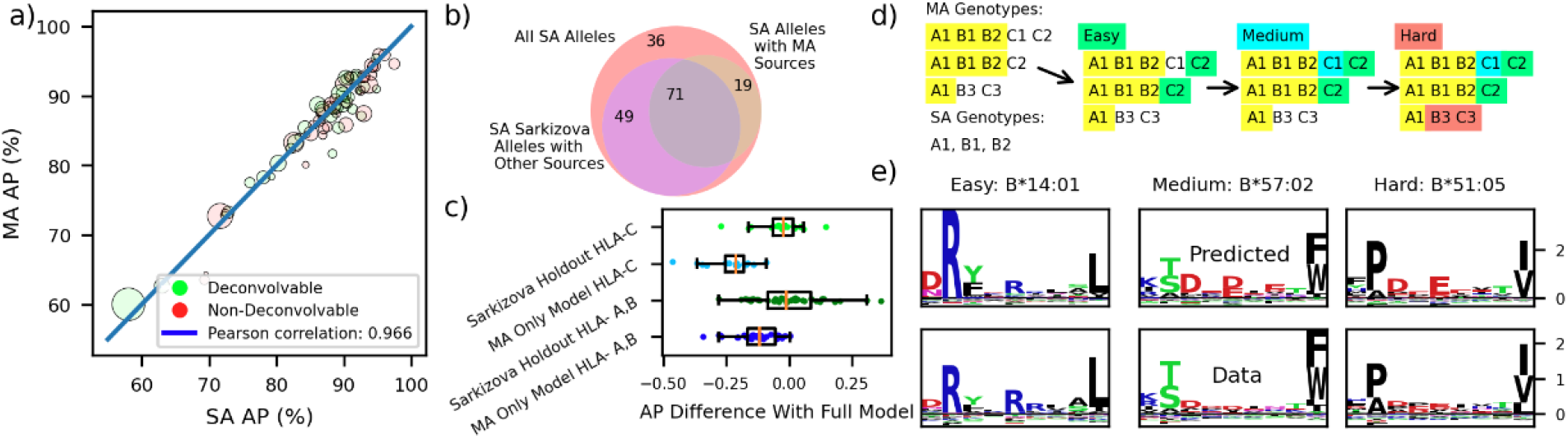
HLApollo performs de novo deconvolution on MA data a) depicts a scatter plot of per allele AP obtained from a model trained using only synthetic MA data, and a model trained with just the SA version of the dataset. Both models are evaluated on the same SA dataset. Alleles determined to be deconvolvable and non-deconvolvable according to the Bassani-Sternberg definition are plotted in green and red, respectively. A Pearson’s correlation of 96.6% is observed between the APs of each model. b) depicts a Venn diagram of the various sources of SA data available to train a model, all SA alleles (red), all SA alleles from Sarkizova et al. that also have other study sources (purple), and SA alleles with MA sources (brown-orange). The grey-blue circle represents SA alleles which exist in Sarkizova et al. and have other data sources, and which also have MA sources, this is used for evaluation in c). c) depicts boxplots of the AP drop relative to the full model evaluated on SA Sarkizova alleles with MA sources. The models evaluated are: 1) Sarkizova holdout (a model trained without data from Sarkizova et al.) and 2) MA only model (a model trained with only MA data). Upper (light green) depicts the performance drop of just HLA-C alleles of the Sarkizova holdout, upper middle (light blue) depicts the performance drop of just HLA-C alleles of the MA only model, lower middle (green) depicts the performance drop of the Sarkizova holdout on HLA-A,B, and lower (blue) depicts the performance drop of the MA only model on HLA-A,B. d) depicts a diagram illustrating our method for allele deconvolvability difficulty, where yellow, green, blue, and red indicate trivial, easy, medium, and hard, respectively. e) depicts motif plots for B*14:01, B*57:02, and B*51:05, which are easy, medium and hard to deconvolve, respectively, predicted by (SA and MA trained) HLApollo (above), and observed (below). HLApollo achieves and AP of 94%, 83%, and 82% for B*14:01, B*57:02, and B*51:05, respectively.

While our deconvolution strategy works well in synthetic data, we also must see if systematic problems with certain alleles (e.g. low presentation of peptides by HLA-C) reduce deconvolution performance in practice in real data. One challenge of real data is that SA samples used for evaluation are also study holdouts. In order to disentangle performance decline caused by deconvolution and generalization to new studies, we create a dataset based on BM1, but with all data from Sarkizova et al.^28^ held out and train a new model. Fig. 5c depicts the AP drop (relative to the full model) of the MA only model and Sarkizova holdout on the 71 SA alleles from Sarkizova et al. that exist in the multiallelic data, and have SA alleles from sources other than Sarkizova (Fig. 5b, BM7). A median AP drop of 1.9% was seen for Sarkizova Holdout, indicating that for some alleles new sample-specific sub-motifs are found inside of the Sarkizova dataset. The MA-only model experienced a more significant median AP drop, of 15.3%. This drop is driven in large part by HLA-C, which declines by 21.1%, indicating that deconvolving HLA-C specificity from MA data is particularly challenging.

We further evaluated how deconvolution ‘difficulty’ impacted HLApollo’s ability to recognize patterns from individual alleles observed in MA genotypes. We propose four levels of difficulty for deconvolution: trivial, easy, medium, and hard, based on their combinatorial uniqueness to MA samples. The definitions are depicted diagramatically in Fig. 5d (see methods). Fig. 5e depicts predicted (MA+SA trained) HLApollo motifs (upper) and observed motifs (lower) from easy, medium, and hard deconvolvable alleles. In order to obtain observed motifs for these alleles, we engineered SA cell lines and obtained ligandomes (see methods for more details), but did not include them in our training dataset. One may observe an improving trend of predicted motifs from easy to hard alleles, as expected. Yet, even for the hard allele, B*51:05, HLApollo predicts a reasonable motif and achieves an AP of 82%, indicating good deconvolution performance.

### Pan-Allelic Generalization

In a clinical setting, full coverage of a subject’s HLA-I genotype by pMHC-I presentation models is desired. Therefore it is critical to have a pan-allelic model with good generalization for untrained alleles. Yet no method currently exists to determine if a model will have good performance on a particular untrained allele. To validate HLApollo’s pan-allelic generalization, we first observe the degradation of AP for alleles for a model trained without them in its training dataset (Fig. 6a) (see Methods). While degradation is observed, the median AP drop is 9.6%, 7% worse than the study holdouts compared in Fig. 5c, but still much better than other ablations and comparator models (Fig. 2).

**Figure 6.**
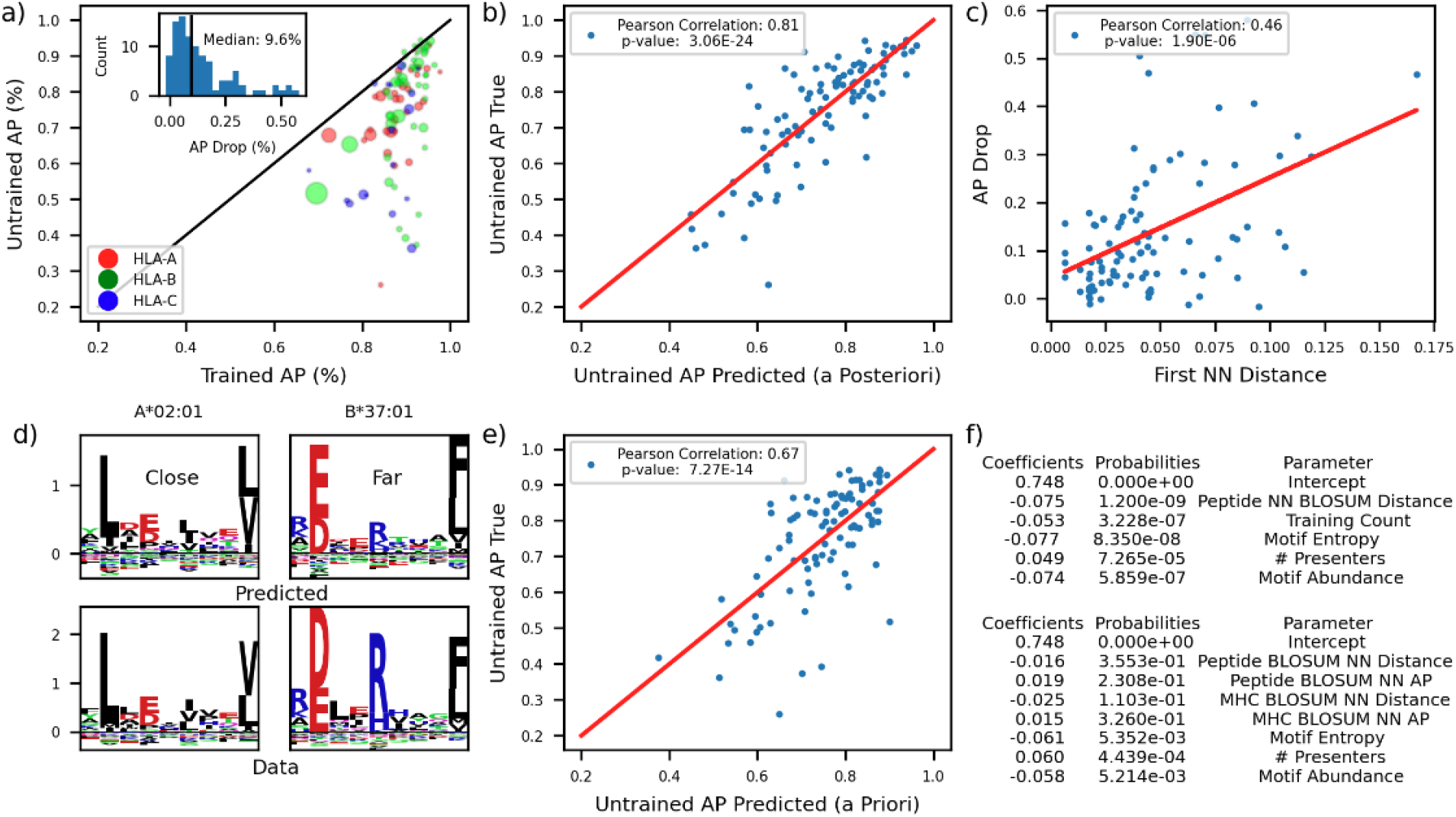
Performance on untrained alleles is predictable a) depicts a scatter plot of trained vs untrained AP for all the single allelic datasets used in training, bubble size indicates number of peptides from the corresponding allele. Inset depicts a histogram of AP drop from full HLApollo b) depicts the correlation between the actual untrained AP, and the predicted untrained AP by the a posteriori untrained AP multilinear regression model. c) depicts the AP drop between trained and untrained models vs MHC NN BLOSUM distance for HLApollo. d) depicts the observed (lower) and untrained HLApollo predicted (upper) motifs for A*02:01 (left) and B*37:01 (right). e) depicts the correlation between the actual untrained AP, and the predicted untrained AP by the a priori untrained AP multilinear regression model. f) depicts the coefficients and probabilities of the parameters used in the a posteriori (upper) and a priori (lower) untrained AP models.

We follow the lead of HLAthena^28^, producing a linear regression model to model AP from attributes of the dataset, but here predicting the out-of-training (OOT) performance. We find this simple model was predictive of untrained AP (Fig. 6b). The parameters used, their coefficients, and p values are shown in the upper half of Fig. 6f (see Methods for in depth description of the parameters). Unsurprisingly, 9mer motif entropy was found to have the largest impact, followed closely by motif abundance and the BLOSUM distance of the OOT 9mer motif’s nearest neighbor (NN) in the trained dataset. The latter feature’s importance indicates that HLApollo reuses representations that it has built for other alleles, a key feature of pan-allelic generalization.

While the model depicted in Fig. 6b is useful for understanding pan-allelic generalization, it has limited clinical applicability because many of the important features are not known a priori. To this end we investigate a feature which is available a priori, the MHC NN BLOSUM distance (Fig. 6c). One may observe modest correlation between the drop in trained vs untrained AP (AP drop) and this metric (Fig. 6c). We observed the untrained (upper) and observed (lower) motifs for the alleles with the closest (left) and furthest (right) MHC NN BLOSUM distance (Fig. 6d). Good agreement for the close allele’s untrained motif and the observed motif is observed, but the distant allele’s untrained motif underrates the importance of the unusual anchor residue at position 5.

Next, we propose a clinically useful, a priori model for predicting untrained AP (Fig. 6e) with MHC NN BLOSUM distance and several other parameters (depicted in the lower half of Fig. 5f). Most of these parameters are similar to those in the a posteriori model, but using (untrained) model predictions on random peptides to extract the parameters. This model achieves reasonable predictive value (Pearson’s correlation of 67%), and so we seek to use it to evaluate if HLApollo predictions should be considered useful for a particular untrained allele. To be considered ‘covered’ by HLApollo we demand a predicted untrained AP of at least that of the worst performer amongst the trained alleles, 67.8%, plus one standard deviation of error in the predictive model, a total of 76.7%. For a comprehensive estimate of genotype ethnic coverage we queried allelefrequencies.net for allele frequencies from only gold-standard datasets with at least 1000 samples (October 6 2021) and evaluated coverage on the 2627 alleles identified^48^. Generalization accounts for HLApollo coverage on 791 alleles, with particularly large gains in genotype coverage amongst several ethnicities, including Asian and Hispanic, shown in Table 1.

**Table 1.** The full genotype coverage (HLA-A1 * HLA-A2 * HLA-B1 * HLA-B2 * HLA-C1 * HLA-C2, i.e. all 6 alleles are available in the training data or are expected to pass a threshold AP value of 76.7% for HLApollo) for various ethnicities. The center column shows the coverage using only alleles found in our training dataset. The right column shows the coverage when pan-allelic generalization is accounted for.

| Ethnicity | Trained Genotype Coverage | Generalized Genotype Coverage |
| --- | --- | --- |
| Amerindian | 82.9% | 86.5% |
| Arab | 86.9% | 87.4% |
| Asian | 79.7% | 84.2% |
| Austronesian | 47.6% | 67.0% |
| African | 89.2% | 90.6% |
| European | 93.8% | 95.1% |
| Hispanic | 70.0% | 80.5% |
| Mestizo | 47.7% | 66.8% |
| Polynesian | 50.1% | 70.5% |

## Discussion

With the recent surge in the availability of large-scale immunopeptidomics data, several deep learning approaches have been applied to the problem of predicting peptide presentation by MHC-I, including convolutional neural networks^34^, feed forward neural networks^28^, mixture models^48^, LSTM^49^, etc. The inductive bias of sequence-based neural networks like transformers and recurrent neural networks is well suited for the problem, as distant amino acids can impact one another’s representation. In this study, we implement a highly accurate, transformer-based approach for pan-allelic peptide presentation prediction by MHC-I, HLApollo. Our approach provides a more generalized representation of the peptide presentation problem compared to heuristics usually implemented in other methods. We achieve this by using a pan-allele approach, handling variable peptide lengths by projecting them to a common embedding vector after the transformer layers, and performing inherent deconvolution when multiallelic data is input. We also introduce a negative set switching approach that increases the diversity of the negative space observed during training without overfitting to it, and substantially adds to the performance, enabling a large model that can capture pan-allelic generalization. Our *bonafide* deconvolution strategy also enables the best use of multi-allelic data to date, achieving the smallest gap between SA-only and MA-only performance on BM1 of the models we evaluated, indicating superior deconvolution capability. Unlike the recent transformer approach, TransPHLA, we do not concatenate all token representations and send them through fully connected layers, as this reintroduces issues with padding (e.g. representation of padded tokens are processed by the model) and leaves only 1% of model parameters in the transformer encoder (which we found led to poor performance on our test dataset).

While other approaches normalize the ranking of different alleles by calculating the % rank of a given peptide with respect to each allele, we opt against this strategy. From Supplementary Fig. 5, one may observe significantly lower HLApollo logit scores for alleles from HLA-C than HLA-A and HLA-B, we believe this to represent biological differences in the presentation rates between genes, and so avoid normalization of scores for different alleles. Further, we do not use binding affinity data, which is allele-specific as a different reference peptide is used for each allele in binding assays. We also constructed only a single pan-allele model as against multiple allele-specific models. Taken together, a calibration step is not needed in our approach, thus avoiding yet another heuristic. However, we note that inherent allelic biases because of antibody pulldowns might still be present, especially in MA data.

Both NetMHCpan4.1-EL and our ensembled feed forward neural network fell significantly short of HLApollo (Fig. 2) demonstrating the importance of using a sequence-based neural network over adhoc anchor placement conserving padding strategies. This was shown despite important biases that favor NetMHCpan4.1-EL. 34.9 % of our positive BM1 test set evaluation data appears in NetMHCpan4.1-EL’s training data (MixMHCpred2.2 is also trained on 43.24% of our test set). NetMHCpan4.1-EL and NetMHCpan4.1-BA (NetMHCpan4.1 BA is the p-MHCI binding affinity predicted by the NetMHCpan4.1 model) have been used pervasively in selecting epitopes for immunogenicity studies, which can potentially bias results in its favor^27^. Additionally, peptides chosen for BA datasets are often enriched for T-cell epitopes, and many studies investigating T cell response use a binding affinity model to select peptides^29^.

Besides modeling peptide presentation purely based on amino acid sequences, we also considered other contributing factors, like expression of the source gene and sequence properties of the source protein. Inclusion of expression information boosted accuracy, primarily by attenuating sequence-based prediction at low expression levels. We argue that protein compartmentalization and other features of source proteins that affect their propensity for peptide presentation are encoded in the protein sequence, and develop a gene-level propensity score, ESM_MHC-I_, by training protein language representations to presentation likelihood. ESM_MHC-I_ also boosts the accuracy of HLApollo, and to a better extent than including manual annotations like Gene Ontology. Surprisingly, the boosts by expression and ESM_MHC-I_ were not synergistic, indicating that ESM_MHC-I_ might have intrinsically encoded some expression biases by protein families. For instance, transcription factors are generally expressed at low levels. Our findings also suggest that HLApollo + ESM_MHC-I_ can be useful in applications where expression information is limited. This method showed performance gains in tissue hold-out presentation data, and also in T cell response data (Fig 4). However, when actual expression information is available, HLApollo + Expr shows superior performance (Fig 4e).

Lastly, pan-allelic modeling is critical for improving patient inclusion in clinical trials, and so it is a primary goal of this work to investigate the generalization of HLApollo to new studies and new alleles. The linear regression model in Fig. 5e enables the prediction of model performance on untrained alleles using only a priori knowledge gathered from either the allele pseudo sequence or the model’s preconceived notions of the allele. Using confident generalizers benefits people of color, who are unfortunately underrepresented in currently available ligandome data. This method also enables the selection of alleles for acquiring ligandomes by prioritizing predicted low performers with large population frequencies.

## Methods

### Generation of Single-Allelic cell lines and cell culture

HMy2.CIR (C1R) cells were chosen due to their ease of culture and lack of endogenous HLA-A and HLA-B protein expression. HLA-C was further disrupted using CRISPR/Cas9, resulting in an HLA Class I null parental cell line. Stable mono-allelic cell lines were generated through transduction of the Class I null parental line with lentiviral constructs expressing the HLA allele of interest. Enrichment of HLA-expressing populations was performed by magnetic bead based isolation with a pan-Class I antibody (W6/32). Population purity was confirmed by flow cytometry using the same antibody following expansion. The HLA genotype was validated by confirmation of an incorporated sequence barcode before further processing (*reference Gurung, Heidersbach, Darwish et al currently unpublished*).

### pMHC-I immunopurification, LC-MS/MS

Immunoprecipitation was performed on 500 x 10^6^ cells for each sample. The cells were lysed in 0.25 % Sodium deoxycholate, 200 μM iodoacetamide, 1% N-Octyl-β-D-thioglucoside, 1 mM EDTA,1 protease inhibitor tablet per 10 mL of DPBS buffer and immunoprecipitated with pan HLA-A, B, C antibody (clone W6/32) that was immobilized and covalently linked to Protein A cartridges (Agilent G5496-60000). Peptides were acid eluted from antibody bound ProteinA cartridges using 0.1M acetic acid/0.1%TFA followed by desalting with C18 solid-phase extraction (SPE). The eluted peptides were injected onto a 75um analytical column packed with Luna C18 resin (Phenomenex) and separated at a flow rate of 350nL/min. The LC eluent was directed into an Orbitrap Fusion Lumos Tribrid mass spectrometer (Thermo Fisher Scientific) equipped with a nano-electrospray ionization source with spray voltage set at 1,900 V. Mass spectral data were acquired using Orbitrap MS scans (R = 60,000 at m/z 400), followed by Orbitrap MS/MS scans (R = 15,000 at m/z 400) subjected to CID and EThCD fragmentation.

### MS data analysis and peptide identification

The tandem mass spectral data were searched using PEAKS Studio (v10.5)^50^ against the Swiss-Prot human database (downloaded 01 Jan 2020) with no enzyme specificity, peptide mass tolerance at 10ppm, fragment mass tolerance at 0.02Da, variable modifications on Oxidation (M) and Deamidation (NQ). Search results were filtered to an estimated peptide false discovery rate of 1%.

### Ligandome data processing

#### Positive Elution Data

Positive peptides originally derive from searching spectra generated by liquid chromatography–tandem mass spectrometry experiments on eluted MHC ligands. Data was collected from 23 published studies (Supplementary Table 2) along with IEDB (originally from^34^) and deposited in a custom sqlite database, mhcDB.

For training and evaluating presentation, we filtered a total of 142,298 positive peptides for a variety of reasons. Peptides were mapped to translated transcripts from Gencode 27, Basic set (GRCh38.p10)^51^ for the purposes of unambiguously modeling cleavage and source gene mRNA expression. Peptides that mapped to multiple genes were dropped from training and test sets, to enable unambiguous training of peptide presentation based on gene expression (54,056 peptides). Similarly, peptides that mapped to multiple isoforms of the same gene were dropped if they had different flanking sequences of 10 amino acids, to enable unambiguous encoding of peptide processing information (42,540 peptides) (Fig. 1c). Moreover, peptides that did not unambiguously map to the human proteome were discarded from presentation benchmark sets (36,553 peptides). Additionally, we did not consider peptides with post-translational modifications (24,341 peptides) or that fell outside the length range of 8-14 amino acids (29,566 peptides). An additional 21,816 positive peptides were excluded from presentation training and evaluation as these were segregated for future internal use as a test set.

#### Negative Data

Since experimental negative peptides cannot be obtained from current approaches, we use the non-presented parts of the human proteome to sample negative peptides. Notably, a small fraction of these computationally identified negatives might be presentable, but were not identified in the current ligandome data. These are ‘gold std false negatives’, and presumably they constitute a small fraction that will not affect machine learning significantly. First, we sampled negative peptides without replacement, per genotype, across the human proteome with a uniform probability of peptide lengths, 8-14. For each single-allelic genotype, positive peptides from other multiallelic genotypes with a matching pseudo sequence were excluded from the negative set. For each multi-allelic genotype, positive peptides from matching single-allelic genotypes with a matching pseudo sequence were also excluded from the negative set.

### BM1, BM3, Sarkizova Holdout, BM7, K-folds

For BM1 (presentation evaluation) the positive elution benchmark dataset was first constructed, per genotype, by randomly selecting 90% of available {peptide,genotype} tuples for training and 10% for testing after filtration (see above). Next, the negative training benchmark dataset was constructed, per genotype, at a 1:1 negative:positive ratio for 520 mutually exclusive sets of {peptide,flank,genotype} tuples in the training split and 4999:1 negative:positive ratio for a single set in the test split. We subsetted this to a ratio of 99:1 for evaluation of HLApollo (Fig. 2), and kept the 4999:1 ratio for evaluation of HLApollo + Expression (Fig. 3). Finally, to eliminate any possible test set leakage, we enumerated all possible {el,peptide,pseudosequence} tuples in the train and the test splits, took their intersection, and then dropped, from the test data, any {el,peptide,genotype} tuples that mapped to this intersection.

The BM3 training dataset for evaluating tissue holdouts was constructed from BM1 by removing any genotypes that matched those derived from the Schuster, Loffler, or Pyke studies according to our mhcDB. Then the dataset was rebalanced at a 1:1 negative:positive ratio per genotype. The BM3 test dataset was taken directly from Supplementary Data 1c from^27^. To account for missing flanking sequences from this test set, peptides were remapped to the human proteome (GRCh38.p10) and flanks of length 10 and 30 were used to predict HLApollo and HLAthena MSiC, respectively. In cases where a single peptide mapped to multiple flanking sequences, HLApollo and HLAthena predictions were averaged. In cases where a single peptide did not map to the proteome, flanks were not considered in the HLApollo prediction and HLAthena MSI score was used. Finally, to eliminate any possible test set leakage, we enumerated all possible {el,peptide,pseudosequence} tuples in the train and the test splits, took their intersection, and then dropped any {el,peptide,genotype} tuples from the test set that mapped to this intersection.

The Sarkizova Holdout training set was constructed by excluding all {peptide,genotype} tuples derived from the Sarkizova study from the BM1 training dataset. Similarly to BM1, we maintained a 1:1 negative:positive ratio in the training split across 520 sets of negative {peptide,flank,genotype} tuples. The Sarkizova Holdout test set was generated as the subset of the BM1 test set derived from the Sarkizova study and a 99:1 ratio was maintained in the test set. The Sarkizova Holdout MA model was trained only on the MA subset of the Sarkizova Holdout training set.

The K-folds datasets were generated similarly to BM1. First, the rows of the BM1 positive data were randomly shuffled. Next, the shuffled dataset was evenly split into 5 sets. For each of the 5 sets, 1 was used as a test set and the remaining 4 were used for training. For each of the 5 sets, negative train and test data was generated similarly to BM1. For each of the 5 sets, test set leakage was removed similarly to BM1, except using 50 negative ensembles instead of 520.

### BM6 (Deconvolution Simulation)

While controlling for peptide count and pairwise motif similarity, we restricted half of the alleles to belong to unique combinations of synthetic samples. We term these as ‘deconvolvable’ alleles, as presumably their motifs would be expected to be present in the unique combination of samples, hence identifying them. The other set of alleles were assigned to non-unique sets of synthetic samples (hence termed ‘non-deconvolvable alleles’)

Alleles with at least 1000 unique peptides (72 alleles) were first selected to mitigate performance bias due to lack of training data. To ensure even distribution of peptide counts across alleles, each allele contributes exactly 500 randomly selected peptides to a synthetic sample. The SA training data was constructed from this dataset, containing 140,530 unique {allele, peptide} tuples.

For MA training data, similar to the biological scenario, a minimum of 1 and maximum of 2 alleles per gene (A, B, C) were necessary to define a synthetic sample. To attenuate the potential impact of pairwise allele similarity on deconvolution performance, pairwise allele similarities informed constraints on which pairs of alleles were allowed to belong together in synthetic samples. Specifically, distance was defined as Jensen Shannon Divergence between 9-mer peptides in the training data of the selected alleles and a threshold of 1.23 was determined as the minimum distance allowed between two alleles (based on visual inspection of motifs). Any peptides that were duplicated per sample were removed.

To explore model performance in the context of principles of deconvolvability of allele-specific peptides based on co-occurrence and exclusion of alleles across samples, roughly half (36/72 alleles) were randomly forced to always occur in pairs when assigned to synthetic samples (‘non-deconvolvable’). The remaining alleles were assigned randomly to synthetic samples, each occurring in a unique set (‘deconvolvable’).

To ensure a dataset of realistic size, 195 synthetic samples were generated, yielding 514,029 total unique genotype-peptide tuples encompassing 107,362 unique peptides.

To ensure no overlap of positive and negative peptides when combining alleles, any overlapping positive/negative SA peptides were removed from consideration of the negative peptides for a given synthetic sample.

We define the 4 categories of deconvolution difficulty as follows: Trivial deconvolvable alleles found in MA genotypes are those for which SA data exists in our dataset. Easy deconvolvable alleles are those for which SA data is absent, but all other alleles within the genotype in MA data are trivially deconvolvable, thus effectively making the remaining allele deconvolvable. Medium deconvolvable alleles are those for which all other alleles within the genotype are either trivial or easy deconvolvable alleles, leaving the medium allele to be deconvolvable. Hard deconvolvable alleles are those for which no combination of trivial, easy, medium (or iterations beyond medium, where medium alleles are used to deconvolve remaining alleles) alleles can isolate the allele. For comparison with the co-occurence and exclusion definitions of deconvolvability, both medium and hard deconvolvable alleles are considered ‘non-deconvolvable’.

### Gene Expression

#### Data Processing / QC

Fastq files were collected from bulk RNAseq expression profiling studies on specific ligandome samples (e.g. Shraibman_MA_2016), subjects (e.g. cell lines, Schuster_ovarian_2017), and representative healthy tissue atlas specimens. First, RNAseq sample runs were aligned using HTSeqGenie^52,53^.

Sample runs with fewer than 10 million unique/concordant mapped reads were discarded from downstream analysis. Other QC metrics (e.g. percent mapped ribosomal reads, mitochondrial reads, etc.) were visually inspected but no sample runs exceeded recommended QC cutoffs. After initial upstream QC, samples were pseudo-mapped / quantified using Salmon (v1.3.0)^54^ on the Ensembl90 reference transcriptome. This approach allowed us to increase sensitivity via consideration of multi-mapped reads and quantify transcript expression as a function of sequence-specific bias, thereby attenuating potential batch effects.

We also considered such a heterogeneous dataset may contain batch effects due to various technical biases such as paired-end vs. single-end sequencing, strand protocol, read length, and sequencing instrument. We conducted principal component analysis to examine such biases. No principal components obviously separated technical features (shown by visual inspection, binomial/multinomial regression of principal components onto features).

#### BM2 & Expression Quantification

We matched RNAseq expression to immune peptidomics data in BM1 by joining RNAseq assay data (Database table mhcDB::Assay_rnaseq) on one of either: mhcDB::Sample.id, mhcDB::Subject.id, or mhcDB::Tissue_atlas.id - depending on the granularity available. We define the entity RNAseq sample as a unique combination of: mhcDB::Sample.id, mhcDB::Subject.id, mhcDB::Tissue_atlas.id, mhcDB::Genotype_mhc_I_human.id.

We quantified gene expression for a sample as the sum of transcript TPM per RNAseq assay from that sample. Then we averaged gene expression across all the different RNAseq assays, if multiple assays were available, for a given RNAseq sample (defined above).

We construct our expression evaluation dataset independently for each RNAseq sample, and in each case, restrict the universe of possible peptides (+ve and -ve) to 50,000 total peptides. Hence, the ratio of negative to positive peptides may vary between RNAseq samples, but the proportion of positives can be interpreted as the probability of peptide presentation for a given RNAseq sample. More specifically:

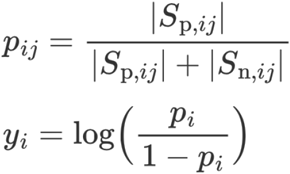

where

*S_P,ij_*: Set of positive peptides in sample *i* in expression bin *j*

*S_n,ij_*: Set of negative peptides in sample *i* in expression bin *j*

*P_ij_*: Probability of presentation in sample *i* in expression bin *j*

*y_ij_*: Empirical enrichment (log odds) for presentation in sample *i*

Note, the total set of peptides considered across the expression bins *J* for a sample *i* is always 50 × 10^3^

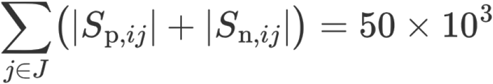

### GO Gene Sets

We used gene sets that correspond to the cellular component (CC) terms from Gene Ontology (GO) ^55^. Specifically, we selected GO CC terms that consist of major subcellular localization based on manual interpretations, and cellular components that have been shown to be associated with MHC-restricted peptides ^14,56^, namely: nucleus (GO ID, GO:0005634), mitochondrion (GO:0005739), proteasome complex (GO:0000502), endoplasmic reticulum (GO:0005783), Golgi apparatus (GO:0005794), cytosol (GO:0005829), cytoskeleton (GO:0005856), cytoplasm (GO:GO:0005737), plasma membrane (GO:0005886), cell junction (GO:0030054), extracellular region (GO:0005576), lysosome (GO:0005764), endosome (GO:0005768), early endosome (GO:0005769), late endosome (GO:0005770), endocytic vesicle (GO:0030139), cytoplasmic vesicle (GO:0031410), vesicle (GO:0031982), recycling endosome (GO:0055037), and secretory vesicle (GO:0099503). Gene annotations found associated with the respective GO CC terms were then obtained from Ensembl release 90 ^57^.

### Motif logos

Positive single-allelic peptides and their (N,C)-terminal flanking sequences were taken from BM1. For motif visualization, positive ligands from BM1 were converted to 8-mers by taking the first 7 amino acids and the last amino acid residue. The background distribution of amino acids and of terminal ends of proteins were computed from the proportions found in the negative peptides (and their flanking sequences) from BM1 (ensembles 1-5, note that the negatives are genotype-specific, so anchor preferences for different alleles get diluted over the space of all negatives). A matrix P_ai_ was computed and used as input to R’s ggseqlogo package or Python’s logomaker package^58,59^ to plot the motif logo.

#### Amino acid (*a*) frequency *p_ai_* at position *i*, averaged across a set of HLA genotypes *G*

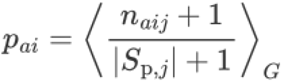

*n_aij_*: Number of occurrences of amino acid *a* at position *i* in positive peptides in genotype *j*.

*S_p,j_*: Set of positive peptides identified for genotype *j*

#### Background frequency *q_a_* of amino acid *a*

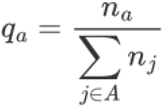

*n_a_*: Number of occurrences of amino acid *a* in all negative peptides and their flanks.

#### KLD for position *i* in a motif logo

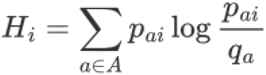

Height of an amino acid *a* at position *i*

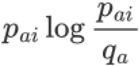

### Model Architectures and Training

All artificial neural network models in this work are implemented in pytorch^60^, and training loops in fastai^61^, on (which includes features such as the default fit one cycle learning rate scheduling) with mixed precision. Binary cross entropy loss and the Adam optimizer^62^ are used in their default settings. HLApollo uses a batch size of 3072 and a learning rate of 0.0002, except for the final transformer encoder layers and fully connected layers, which use a learning rate of 0.0004. Weights are initialized with default Xavier initialization. Single allelic (SA) samples are upweighted in the loss function by a factor of 4. The feed forward neural network (FFNN) uses a learning rate of 0.0006, and batch size of 900. HLApollo is trained for 115 epochs, and FFNN is trained for 50 epochs. HLApollo’s training time on one machine with 8 NVIDIA V100s is 7.7 hours per model. Ensembling is performed by averaging the predictions from each model, with 10 models. The datasets are subsampled to 80% each epoch to increase diversity across ensembles. Manual hyperparameter optimization is performed for each model, due to the training time involved for each model. Negative set switching is implemented by sampling a new negative set from a space of 500 full negative sets (fewer for those depicted in Fig. 2b) per epoch.

HLApollo’s architecture is as follows. Amino acid tokens are embedded by a learned embedding (pytorch’s nn.embedding) to a dimension of 400. Flanking sequences that reach the edge of the protein are given a special token (this token is padded to the length 10 if relevant). As described in the main text, flanking sequences are randomly dropped 50% of the time, and 50% of the time 0-10 amino acids are removed (following a uniform distribution). The n-flanking, peptide, and c-flanking sequences are padded to their maximum lengths (10, 14, and 10 respectively) and concatenated together (now termed the concatenated sequence) and standard transformer positional encoding is performed. The MHC sequences are treated with a separate embedder and positional encoder (same implementation as for the peptide sequence); all layers that process MHC sequences have shared weights for the various MHC sequences in the multiallelic (MA) setting. The concatenated sequence (peptide and flanks), and its padding mask (to prevent unused tokens from impacting other token’s representations), are sent through a transformer encoder layer. All transformer encoder layers used have a dimension of 400, 16 heads, and no dropout. The MHC sequence is separately passed through its own transformer encoder layer. The concatenated sequence and MHC sequence are then further concatenated, for each MHC sequence, and a beginning of sequence (BOS) token (passed through its own embedder and positional encoder) is concatenated to the beginning of each sequence, called the pair sequence. Without gradient accumulation, all pair sequences are passed through the output module: 4 transformer encoder layers. Next, the BOS token is fed through, sequentially, a fully connected (FC) layer (dimension 256), a dropout layer (50%), a swish activation^63^, a second FC layer (dimension 128), a second dropout layer (50%), a second swish activation, and finally a FC layer to output a single logit score. Used in this way, the BOS token causes information from the rest of the sequence to flow into the token, creating a sequence representation with a fixed size, regardless of the sequence length. Finally, the concatenated sequence-MHC pair with the highest logit score is identified, and this pair is run through the output module with gradient accumulation (this is our deconvolution strategy). Removing the gradient accumulation for pairs that are not determined by the model to be the presenting pair is used to prevent gradient noise (and increase speed) that would accumulate if each is allowed to pass through the output with gradient accumulation.

The FFNN uses the Netmhcpan^64^ inspired 9mer mapping commonly used in the literature. 8mers were increased to a length of 9 by adding an X token in between two existing amino acids, resulting in 7 enumerated peptide possibilities. Peptides longer than 9 had (length - 9) consecutive amino acids removed from their sequence, resulting in 9 peptide possibilities. N-flanking and c-flanking sequences were padded with X tokens to the maximum flanking sequence length (10). N-flanking sequences, possible peptide sequences, c-flanking sequences, and MHC sequences were concatenated, for each peptide possibility and each MHC sequence (for example a 10mer peptide for a 5-allele sample will have 45 sequences associated with it). These paired sequences were BLOSUM62 encoded^65^, and sent into the FFNN model (input size 1050), without gradient accumulation. The FFNN model is composed of the following sequential layers: a FC layer (dimension 512), relu activation, batch normalization, a second FC layer (dimension 512), a second relu activation, a second batch normalization, and a final FC layer to output to a logit score. The peptide possibility - MHC pair with the largest logit score was then chosen for each input and sent through the model again with gradient accumulation.

The non-pan allelic HLApollo was trained with a similar strategy to MHCNuggets^49^, the largest SA dataset (B*27:05) was used to train HLApollo as described above, then transfer learning was used (learning rate 10% as above) to train the other SA models.

ESM1b protein features were extracted by taking token representations of amino acid residues in a particular protein following the details described in their github^66^. To produce a protein-level feature set we averaged the residue features across the protein, resulting in a 1280 dimensional vector. The ESM_MHC-I_ model is a FFNN model that takes the 1280 dimensional protein vector from ESM1b and maps it to a presentation likelihood using two FC layers (500 and a 50 dimensional), with dropout (50%) applied after each hidden layer. We used a batch size of 1024 and a learning rate of 0.001. The model was trained by taking each peptide from our baseline training dataset, determining the protein from which it was derived, and using its protein features as input.

Logistic regression models used to add expression, gene ontology (GO), and ESM_MHC-I_ features to HLApollo score were implemented in sklearn using default settings, except the maximum iterations allowed were set to 3000. Pseudo counts were added to the TPM from the source gene and it was log transformed.

We calculated the % rank by acquiring predictions on random peptides of length 8-14 paired with alleles in our dataset and mapping the logit score to the % rank on this dataset. This is done across all alleles, as opposed to an allele-specific normalization approach. The dataset’s allele distribution is chosen to match that of our test dataset, and the peptide length distribution is uniform.

### Pan-allelic Generalization Methods

Each allele out of training (OOT) model is trained by removing an allele with SA data, and all MA genotypes that the allele appears in from the training dataset, this is performed for each allele with SA data. Training is performed as described above.

The prior knowledge linear regression model, implemented in sklearn using default settings, is trained using the following (scaled) parameters: 1) The peptide BLOSUM distance to its nearest neighbor (NN) in the training dataset. Amino acids are embedded in the BLOSUM62 space, and averaged position wise across each allele’s peptides, and then these 9 vectors are concatenated to form one vector for each SA allele, thus representing the ‘average’ ligand for each allele. For each held-out allele, the training allele with the minimal distance in this space is found and this distance is used as the peptide BLOSUM distance to its NN. 2) The number of peptides in the SA dataset of interest. 3) The peptide entropy of interest. This is calculated by generating a motif for each 9mer peptide of interest (implemented with logomaker^67^), and summing the entropy over the motif. 4) The # of peptides with HLApollo logit score over 0 on a dataset of 100,000 random peptides. 5) The motif abundance, proposed in Sarkizova et al.^28^ which sums the frequency of each amino acid in a 9-mer motif normalized by the frequencies of the amino acid occuring in the human proteome.

The a priori linear regression model, also implemented in sklearn using default settings, is trained with the following (centered and scaled) parameters, which have all been obtained by getting the appropriate OOT HLApollo predictions on 100,000 random peptides, and considering those with logit score over 0 to be likely presenters: 1) The peptide BLOSUM distance to its NN in the training dataset. 2) The AP of the model on this peptide-based NN dataset. 3) The MHC BLOSUM distance to its nearest neighbor. 4) The AP of the model on this MHC-based NN dataset. 5) The predicted motif entropy. 6) the predicted # of presenters out of 100,000. 7) The predicted motif abundance. Inputs to both the prior knowledge and a priori linear regression models are centered and scaled prior to fitting.

Allele and genotype coverage is assessed by querying allelefrequencies.net for allele frequencies from only gold-standard datasets with at least 1000 samples (October 6 2021) and evaluated coverage on the 2627 alleles identified^48^. Allele coverage is found per ethnicity and is converted to allele coverage by the following equation:

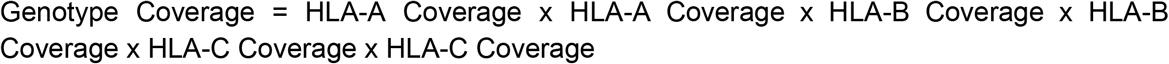

Untrained alleles are considered covered by HLApollo if the a priori model determines that the AP is one standard deviation above the worst performer amongst the trained alleles, 67.8%, a total of 76.7%.

## Supplementary Information

**Supplementary Table 1.** Overview of the modeling capabilities of pMHC-I modeling frameworks. HLApollo is the only modeling framework that has all capabilities.

| Method | Year | BA | Pan<br>allele | Deconvo-<br>lution | Cleav-<br>age | Expr. | GO<br>protein<br>features | ESM<br>protein<br>features | Negative<br>Switching |
| --- | --- | --- | --- | --- | --- | --- | --- | --- | --- |
| HLApollo | 2022 | X | ✓ | bonafide | ✓ | ✓ | ✓ | ✓ | ✓ |
| TransPHLA | 2022 | X | ✓ | X | X | X | X | X | X |
| DeepAttention<br>Pan | 2021 | ✓ | ✓ | X | X | X | X | X | X |
| SHERPA | 2021 | X | ✓ | pseudo | ✓ | ✓ | ✓ | X | X |
| NetMHCpan<br>4.1 | 2020 | ✓ | ✓ | pseudo | X | X | X | X | X |
| MHCFlurry<br>2.0 | 2020 | ✓ | ✓ | X | ✓ | X | X | X | X |
| MHCNuggets | 2020 | ✓ | X | pseudo | X | X | X | X | X |
| HLAthena | 2020 | X | ✓ | X | ✓ | ✓ | ✓ | X | X |
| ACME | 2019 | X | ✓ | X | X | X | X | X | X |
| MHCSeqNet | 2019 | X | ✓ | X | X | X | X | X | X |
| DeepSeqPan | 2019 | ✓ | ✓ | X | X | X | X | X | X |
| EDGE | 2018 | X | X | bonafide | ✓ | ✓ | ✓ | X | X |
| MixMHCpred2<br>.2 | 2017 | X | X | bonafide | X | X | X | X | X |

**Supplementary Figure 1.**
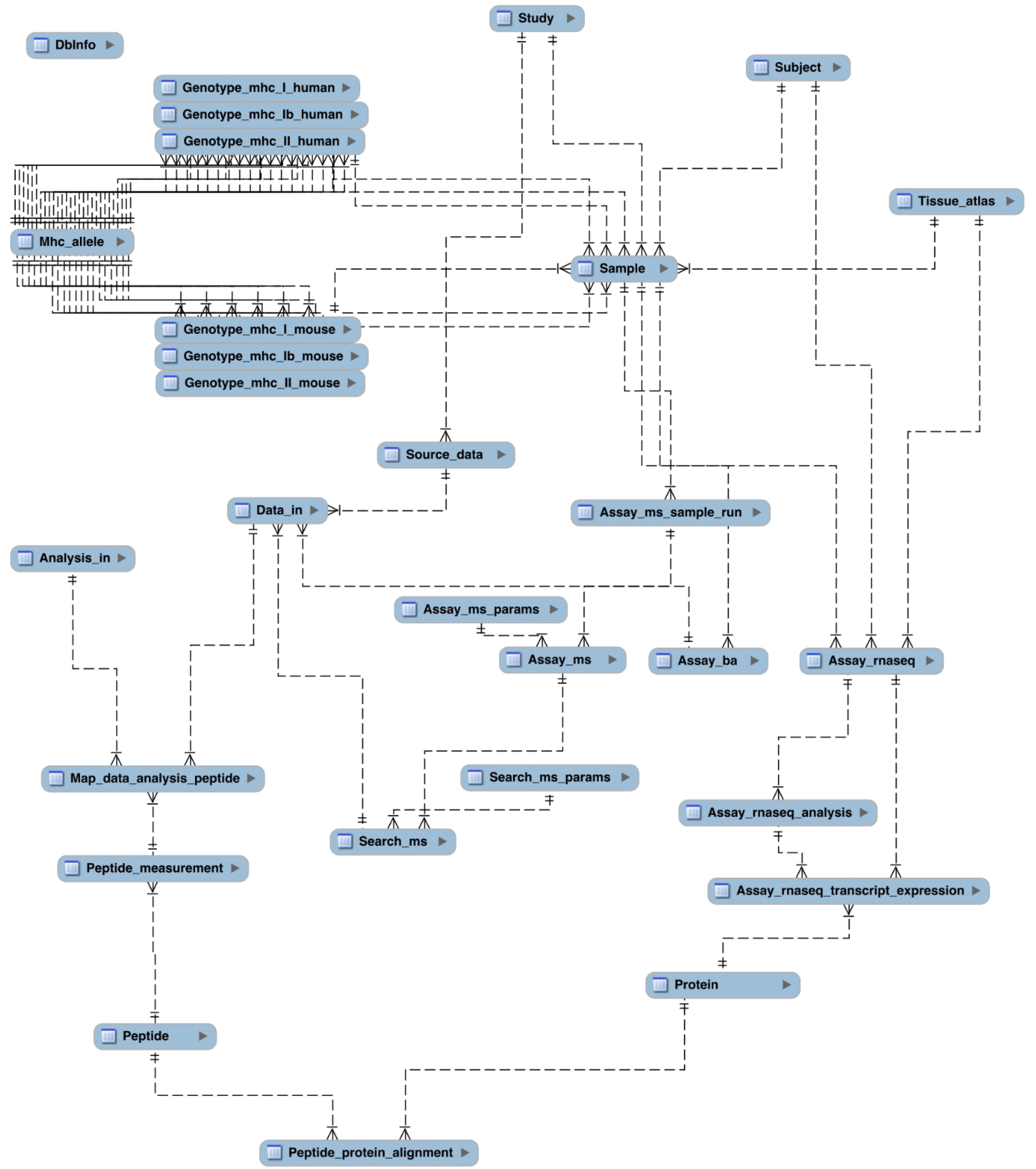
mhcDB schema entity relationship diagram.

**Supplementary Figure 2.**
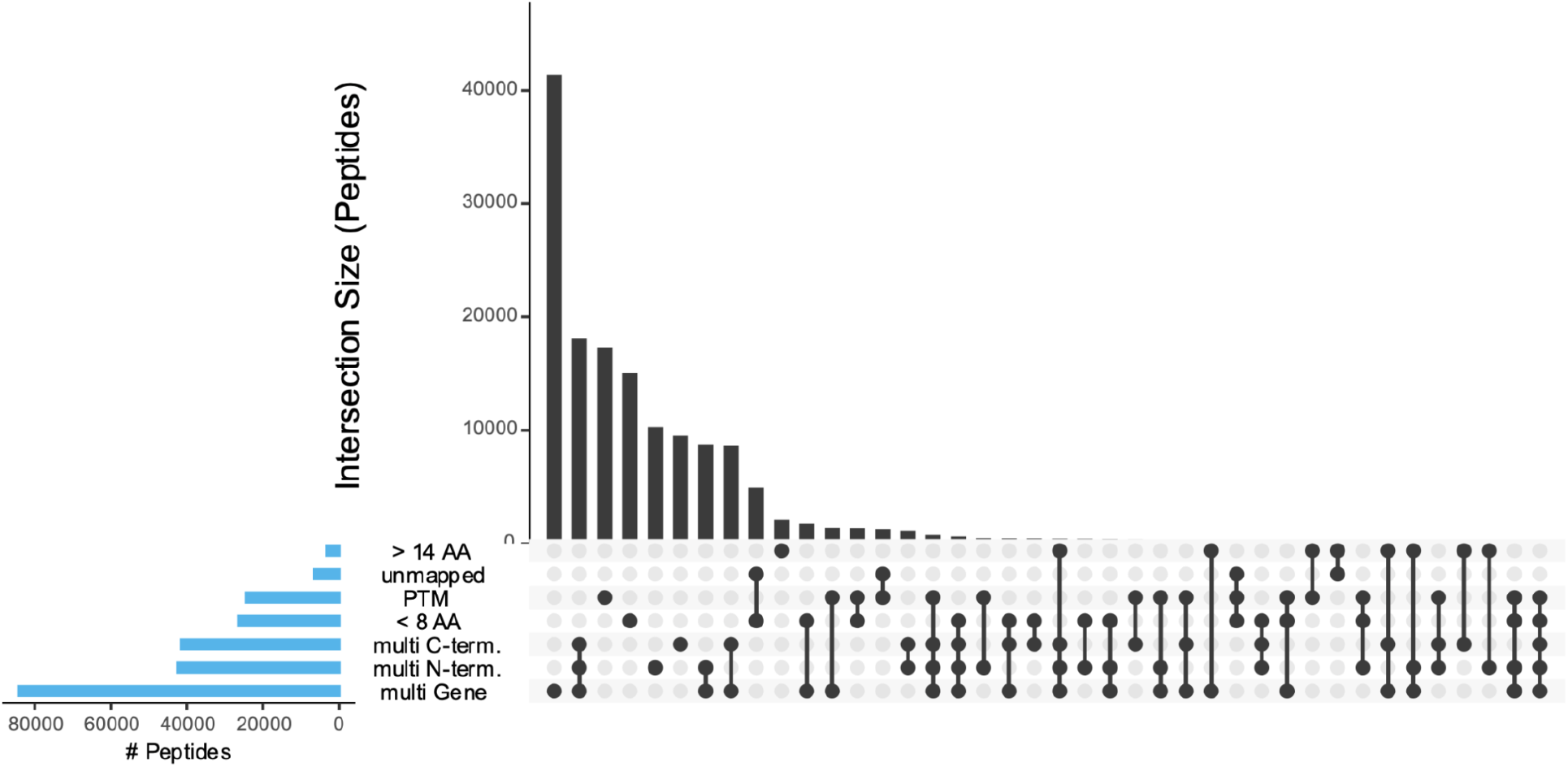
Presented peptides filtered from benchmark datasets.

**Supplementary Figure 3.**
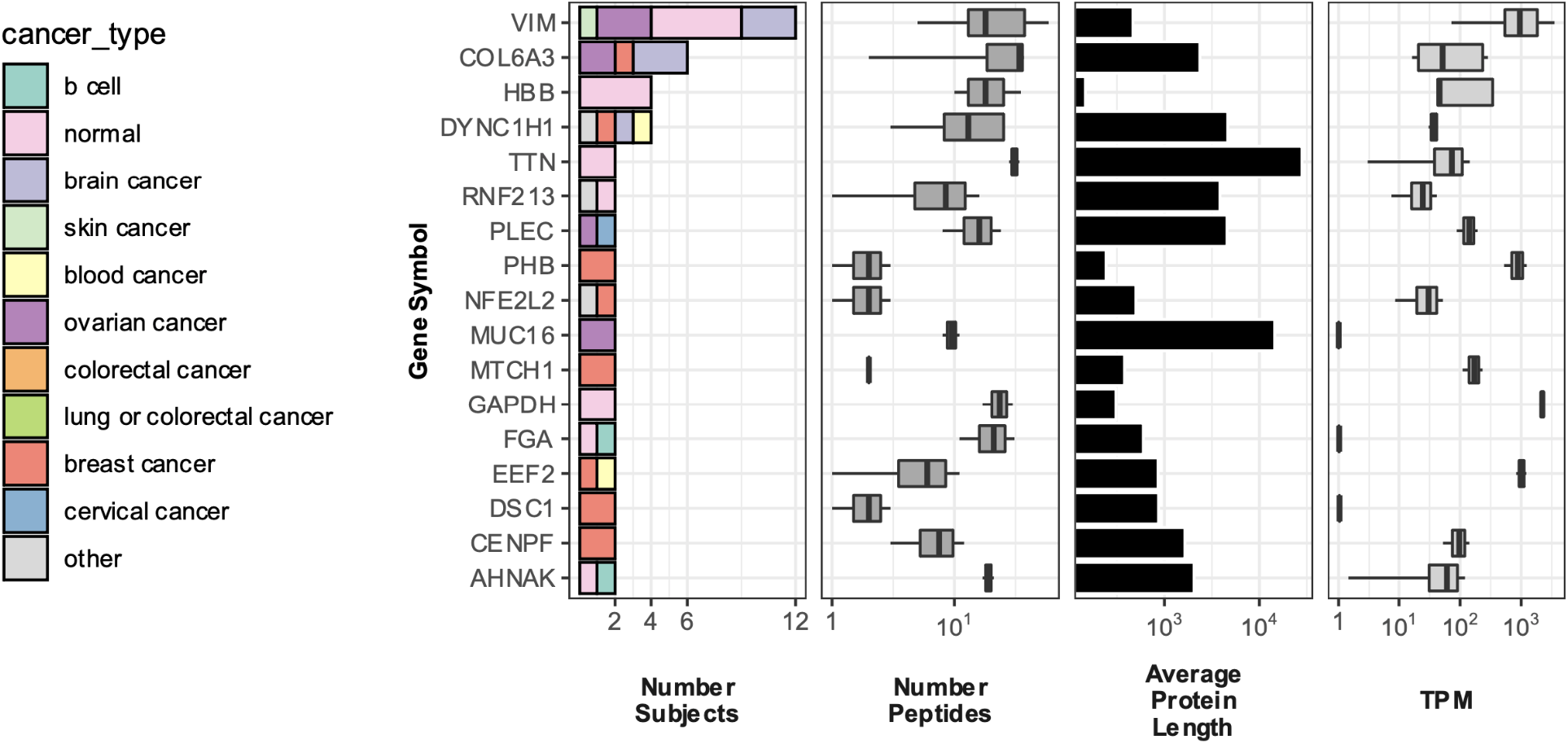
Genes presenting at least one peptide in at least 2 subjects.

**Supplementary Table 2.** 23 studies, including IEDB, used for training and evaluation.

| Study Name | PMID | Description |
| --- | --- | --- |
| iedb_consolidated |  | IEDB curated immunopeptidomics data: 'mhc_ligand_full.csv' |
| sarkizova_mhcl_2020 | 31844290 | mass spectrometry profiling of >185,000 peptides eluted from 95 HLA-A, -B, -C and -G mono-allelic cell lines. (16 alleles previously described in Abelin et. al 2017 Immunity). |
| bassani-sternberg_multiallelic_deconvolution_2017 | 28832583 | Deciphering HLA-I motifs across HLA peptidomes improves neo-antigen predictions and identifies allosteric regulating HLA specificity |
| bassani-sternberg_multiallelic_melanoma_2016 | 27869121 | Direct identification of clinically relevant neoepitopes presented on native human melanoma tissue by mass spectrometry |
| bassani-sternberg_multiallelic_cell-lines_2015 | 25576301 | Mass Spectrometry of Human Leukocyte Antigen Class I Peptidomes Reveals Strong Effects of Protein Abundance and Turnover on Antigen Presentation |
| Mommen_multiallelic_Bcell_2014 | 24616531 | Expanding the detectable HLA peptide repertoire using electron-transfer/higher-energy collision dissociation (ET<sub>h</sub>CD) |
| Gloger_multiallelic_melanoma_2016 | 27600516 | Mass spectrometric analysis of the HLA class I peptidome of melanoma cell lines as a promising tool for the identification of putative tumor-associated HLA epitopes |
| Ritz_multiallelic_cell-lines_2016 | 26992070 | High-sensitivity HLA class I peptidome analysis enables a precise definition of peptide motifs and the identification of peptides from cell lines and patients' sera |
| Shraibman_MA_2016 | 27412690 | HLA peptides derived from tumor antigens induced by inhibition of DNA methylation for development of drug-facilitated immunotherapy |
| Shraibman_GBM_MA_2019 | 31154438 | Identification of Tumor Antigens Among the HLA Peptidomes of Glioblastoma Tumors and Plasma |
| Marcilla_C1R_B2705_2016 | 26811146 | Comparative Analysis of the Endogenous |
|  |  | Peptidomes Displayed by HLA-B*27 and Mamu-B*08: Two MHC Class I Alleles Associated with Elite Control of HIV/SIV Infection |
| Marcilla_C1R_B4002_2014 | 24366607 | Increased Diversity of the HLA-B40 Ligandome by the Presentation of Peptides Phosphorylated at Their Main Anchor Residue |
| Granados_BLCL_2014 | 24714562 | Impact of genomic polymorphisms on the repertoire of human MHC class I-associated peptides |
| Ritz_sHLA_healthyDonor_2017 | 27862975 | Purification of soluble HLA class I complexes from human serum or plasma deliver high quality immuno-peptidomes required for biomarker discovery |
| Alpizar_HLA-B_phosph_2017 | 27920218 | A Molecular Basis for the Presentation of Phosphorylated Peptides by HLA-B Antigens |
| DiMarco_HLA-C_2017 | 28904123 | Unveiling the Peptide Motifs of HLA-C and HLA-G from Naturally Presented Peptides and Generation of Binding Prediction Matrices |
| Chong_Ifng_2018 | 29242379 | High-throughput and Sensitive Immunopeptidomics Platform Reveals Profound Interferon-Mediated Remodeling of the Human Leukocyte Antigen (HLA) Ligandome |
| Hassan_Blcl_2013 | 23481700 | The Human Leukocyte Antigen-presented Ligandome of B Lymphocytes |
| Rozanov_brca_2018 | 29331515 | MHC class I loaded ligands from breast cancer cell lines: A potential HLA-I-typed antigen collection |
| Schuster_ovarian_2017 | 29093164 | The immunopeptidomic landscape of ovarian carcinomas |
| Loffler_crc_2018 | 29789417 | Mapping the HLA Ligandome of Colorectal Cancer Reveals an Imprint of Malignant Cell Transformation |
| Marcu_benign_atlas_2020 | 33858848 | The HLA Ligand Atlas - A resource of natural HLA ligands presented on benign tissues |
| personalis_pyke_mhcl_2021 | 34126241 | 25 monoallelic cell lines + 12 lung/colorectal tumor tissue samples for pan-allele MHC-binding algorithm |

**Supplementary Table 3.** Overview of the datasets used to train and evaluate pMHC-I models. Positive peptide counts are shown for training and test data, while negative peptide strategy is indicated for the test data. All training data used 1:1 +ve:-ve peptide ratio.

| **Dataset** | **Unique Positive Peptide Count (EL=1)** | **Positive Genotype Count (EL=1)** | **Unique Test negatives** | **Model Trained** | **Evaluation** |
| --- | --- | --- | --- | --- | --- |
| BM1 | Train: 259,513<br>Test: 19,740 | Train: 351<br>Test: 235 | +ve : -ve ratio of 1:99 | HLApollo, FFNN, ablations | Naive |
| BM2 | Train: 174,682<br>Test: 11,560 | Train: 181<br>Test: 118 | +ve : -ve ratio of 1:4999 | HLApollo + Expr, HLApollo + ESM, HLApollo + GO | Expression, ESM |
| BM3 | Train: 239,315<br>Test: 1,941 | Train: 289<br>Test: 31 | no negatives | HLApollo (BM3) | Tissue Holdout |
| BM4 | Test: 127 | Test: 34 | 3115 peptides |  | Immunogenicity |
| BM5 | Test (full): 41<br>Test (expr): 33 | Test (full): 12<br>Test (expr): 11 | 867 peptides (full)<br>671 peptides (expr) |  | Immunogenicity |
| BM6 | Train: 107,362<br>Test: 11,757 | Train: 195<br>Test: 72 | +ve : -ve ratio of 1:99 | HLApollo (BM6) | Deconvolution (simulated) |
| Sarkizova holdout dataset | Train: 233,513 | Train: 309 | (train only) | HLApollo (Sarkizova holdout) | Deconvolution (real) |
| MA only dataset | Train: 107,362 | Train: 195 | (train only) | HLApollo (MA only) | Deconvolution (real) |
| BM7 | Test: 10033 | Test: 71 |  |  | Deconvolution (real) |

**Supplementary Table 4.** Ligandome data summary after filtration (Supplementary Figure 2) and before removing peptides overlapping with our private evaluation dataset. The following datasets are a subset of this data: BM1, BM2, BM3, BM6, Sarkizova Holdout, MA only, BM7

| genotype | pMHC-I tuples | alleles | peptides |
| --- | --- | --- | --- |
| multi-allelic | 678,568 | 121 | 203,233 |
| single-allelic | 275,125 | 137 | 180,716 |
| Total | 953,693 | 171 | 305,646 |

**Supplementary Figure 4.**
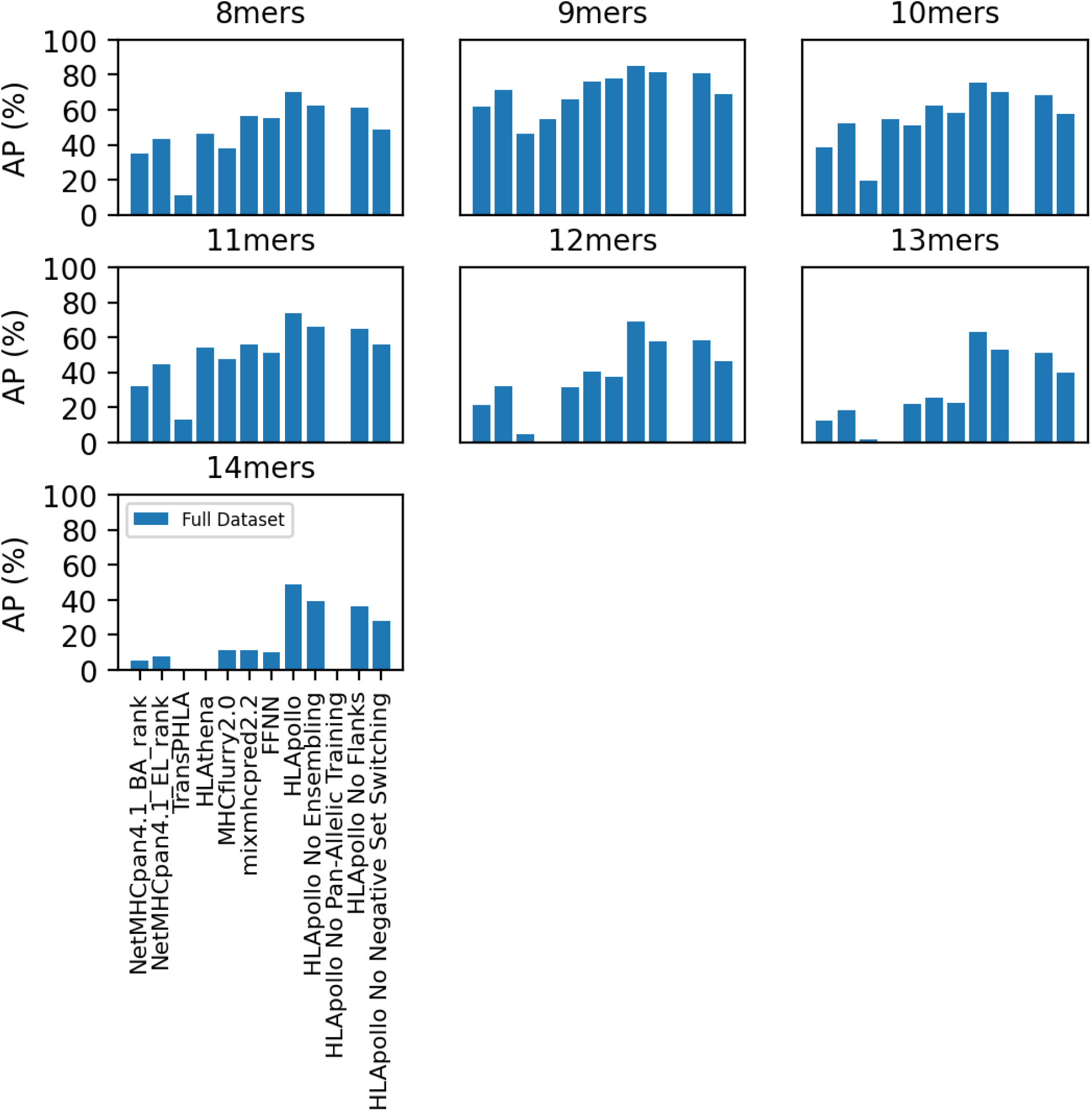
nmer-specific performance of HLApollo(full, ensembled model) on presentation test data(dataset BM1), compared with NetMHCpan4.1, TranspHLA, HLAthena (XXX), MHCflurry2.0, MixMHCpred2.2, and several individual ablations of HLApollo. Performance improvement was seen across all peptide lengths compared to other models and HLApollo ablations. Even for 8-mers, inclusion of pan-allele training improved performance, unlike previously reported^28^.

**Supplementary Figure 5.**
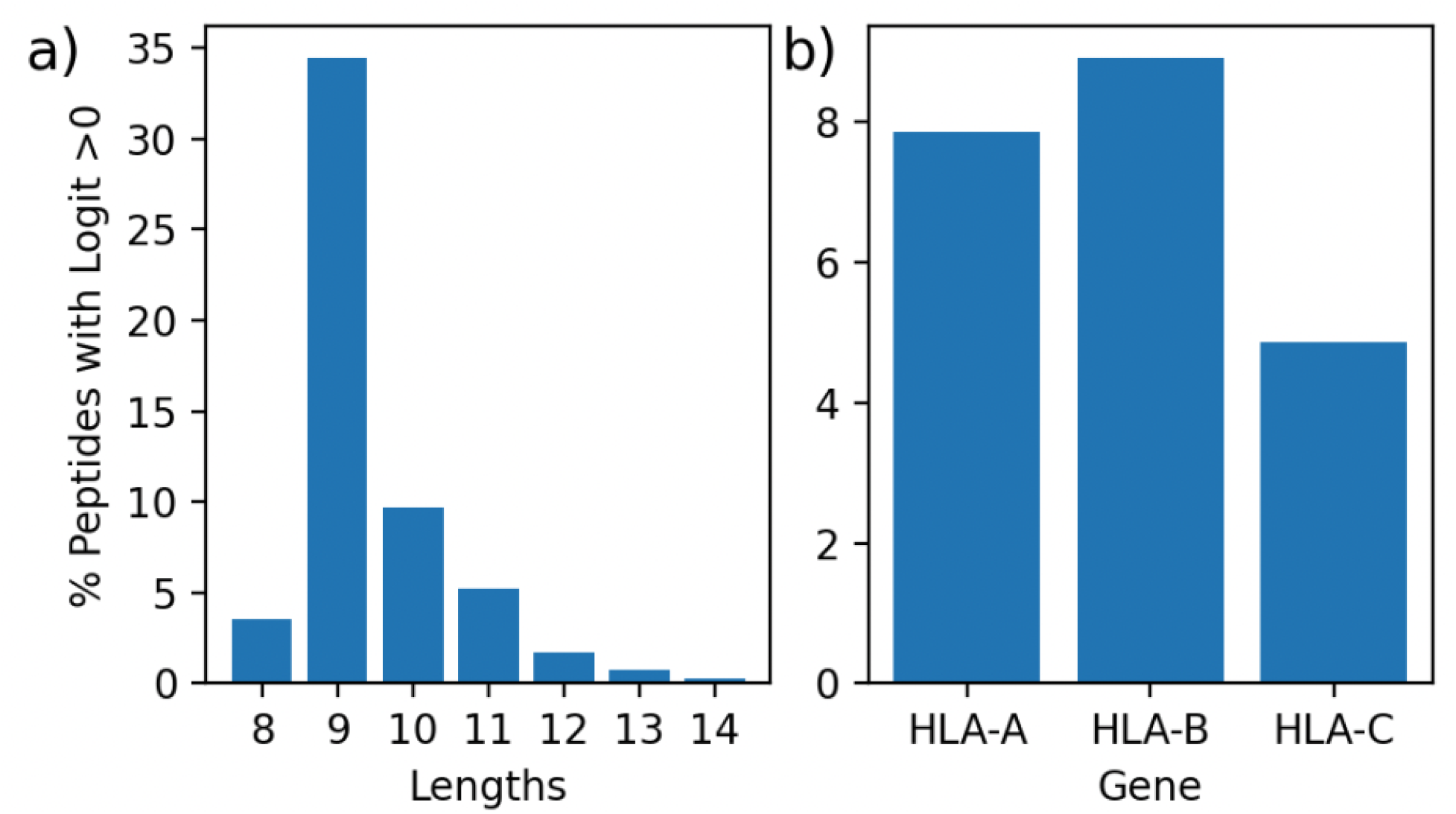
a) depicts the % of random peptides that HLApollo assigns a logit score above 0, dependent on peptide length. b) depicts the % of random peptides that HLApollo assigns a logit score above 0, dependent on the HLA gene.

## References

1. Rizvi, N. A. et al. Cancer immunology. Mutational landscape determines sensitivity to PD-1 blockade in non-small cell lung cancer. Science 348, 124–8 (2015).

2. Rooij, N. van et al. Tumor exome analysis reveals neoantigen-specific T-cell reactivity in an ipilimumab-responsive melanoma. J Clin Oncol 31, e439–42 (2013).

3. Yadav, M. et al. Predicting immunogenic tumour mutations by combining mass spectrometry and exome sequencing. Nature 515, 572–6 (2014).

4. Hugo, W. et al. Genomic and Transcriptomic Features of Response to Anti-PD-1 Therapy in Metastatic Melanoma. Cell 168, 542 (2017).

5. Gubin, M. M. et al. Checkpoint blockade cancer immunotherapy targets tumour-specific mutant antigens. Nature 515, 577–81 (2014).

6. Sahin, U. et al. Personalized RNA mutanome vaccines mobilize poly-specific therapeutic immunity against cancer. Nature 547, 222–226 (2017).

7. Sahin, U. et al. An RNA vaccine drives immunity in checkpoint-inhibitor-treated melanoma. Nature 585, 107–112 (2020).

8. Awad, M. M. et al. Personalized neoantigen vaccine NEO-PV-01 with chemotherapy and *anti-PD-1 as first-line treatment for non-squamous non-small cell lung cancer*. Cancer Cell (2022) doi:10.1016/j.ccell.2022.08.003.

9. Palmer, C. D. et al. Individualized, heterologous chimpanzee adenovirus and self-amplifying mRNA neoantigen vaccine for advanced metastatic solid tumors: phase 1 trial interim results. Nature Medicine 28, 1619–1629 (2022).

10. Jhunjhunwala, S., Hammer, C. & Delamarre, L. Antigen presentation in cancer: insights into tumour immunogenicity and immune evasion. Nat Rev Cancer 1–15 (2021) doi:10.1038/s41568-021-00339-z.

11. Rock, K. L., Reits, E. & Neefjes, J. Present Yourself! By MHC Class I and MHC Class II Molecules. Trends Immunol 37, 724–737 (2016).

12. Neefjes, J., Jongsma, M. L. M., Paul, P. & Bakke, O. Towards a systems understanding of MHC class I and MHC class II antigen presentation. Nat Rev Immunol 11, 823–836 (2011).

13. Trombetta, E. S. & Mellman, I. CELL BIOLOGY OF ANTIGEN PROCESSING IN VITRO AND IN VIVO. Annu Rev Immunol 23, 975–1028 (2005).

14. Abelin, J. G. et al. Mass Spectrometry Profiling of HLA-Associated Peptidomes in Mono-allelic Cells Enables More Accurate Epitope Prediction. Immunity 46, 315–326 (2017).

15. Hunt, D. F. et al. Characterization of Peptides Bound to the Class I MHC Molecule HLA-A2.1 by Mass Spectrometry. Science 255, 1261–1263 (1992).

16. Rammensee, H.-G., Friede, T. & Stevanović, S. MHC ligands and peptide motifs: first listing. Immunogenetics 41, 178–228 (1995).

17. Vita, R. et al. The immune epitope database (IEDB) 3.0. Nucleic Acids Res 43, D405–D412 (2015).

18. Parker, K. C., Bednarek, M. A. & Coligan, J. E. Scheme for ranking potential HLA-A2 binding *peptides based on independent binding of individual peptide side-chains*. J Immunol Baltim Md 1950 152, 163–75 (1994).

19. Nielsen, M. et al. Reliable prediction of T-cell epitopes using neural networks with novel sequence representations. Protein Sci 12, 1007–1017 (2003).

20. Nielsen, M. et al. NetMHCpan, a Method for Quantitative Predictions of Peptide Binding to Any HLA-A and -B Locus Protein of Known Sequence. Plos One 2, e796 (2007).

21. Karosiene, E., Lundegaard, C., Lund, O. & Nielsen, M. NetMHCcons: a consensus method for the major histocompatibility complex class I predictions. Immunogenetics 64, 177–86 (2012).

22. Vaswani, A. et al. Attention is All you Need. in (NeurIPS 2017, 2017).

23. Brown, T. et al. Language Models are Few-Shot Learners. in NeurlPS 2020 (NeurlPS, 2020).

24. Jumper, J. et al. Highly accurate protein structure prediction with AlphaFold. Nature 596, 583–589 (2021).

25. Reynisson, B., Alvarez, B., Paul, S., Peters, B. & Nielsen, M. NetMHCpan-4.1 and NetMHCIIpan-4.0: improved predictions of MHC antigen presentation by concurrent motif deconvolution and integration of MS MHC eluted ligand data. Nucleic Acids Res 48, W449–W454 (2020).

26. Wells, D. K. et al. Key Parameters of Tumor Epitope Immunogenicity Revealed Through a Consortium Approach Improve Neoantigen Prediction. Cell 183, 818–834.e13 (2020).

27. Pyke, R. M. et al. Precision Neoantigen Discovery Using Large-scale Immunopeptidomes and Composite Modeling of MHC Peptide Presentation. Mol Cell Proteomics 20, 100111 (2021).

28. Sarkizova, S. et al. A large peptidome dataset improves HLA class I epitope prediction across most of the human population. Nat Biotechnol 38, 199–209 (2020).

29. Schmidt, J. et al. Prediction of neo-epitope immunogenicity reveals TCR recognition determinants and provides insight into immunoediting. Cell Reports Medicine 2, 100194 (2021).

30. Shao, W. et al. The SysteMHC Atlas project. Nucleic Acids Res 46, gkx664–(2017).

31. Marcu, A. et al. The HLA Ligand Atlas - A resource of natural HLA ligands presented on benign tissues. Biorxiv 778944 (2020) doi:10.1101/778944.

32. Vita, R. et al. The Immune Epitope Database (IEDB): 2018 update. Nucleic Acids Res 47, D339–D343 (2019).

33. Rammensee, H.-G., Bachmann, J., Emmerich, N. P. N., Bachor, O. A. & Stevanović, S. SYFPEITHI: database for MHC ligands and peptide motifs. Immunogenetics 50, 213–219 (1999).

34. O’Donnell, T. J. et al. MHCflurry: Open-Source Class I MHC Binding Affinity Prediction. Cell Syst 7, 129–132.e4 (2018).

35. Wu, S., Du, Y., Beckford, J. & Alachkar, H. Upregulation of the EMT marker vimentin is associated with poor clinical outcome in acute myeloid leukemia. J Transl Med 16, 170 (2018).

36. Sun, B., Fang, Y., Li, Z., Chen, Z. & Xiang, J. Role of cellular cytoskeleton in epithelial-mesenchymal transition process during cancer progression. Biomed Reports 3, 603–610 (2015).

37. Vaswani, A. et al. Attention Is All You Need. Arxiv (2017).

38. Alvarez, B. et al. NNAlign_MA; MHC Peptidome Deconvolution for Accurate MHC Binding Motif Characterization and Improved T-cell Epitope Predictions. Mol Cell Proteomics 18, 2459–2477 (2019).

39. Bassani-Sternberg, M. & Gfeller, D. Unsupervised HLA Peptidome Deconvolution Improves Ligand Prediction Accuracy and Predicts Cooperative Effects in Peptide–HLA Interactions. J Immunol 197, 2492–2499 (2016).

40. Bulik-Sullivan, B. et al. Deep learning using tumor HLA peptide mass spectrometry datasets improves neoantigen identification. Nat Biotechnol 37, 55–63 (2019).

41. Rives, A. et al. Biological structure and function emerge from scaling unsupervised learning to 250 million protein sequences. Biorxiv 622803 (2020) doi:10.1101/622803.

42. Schuster, H. et al. The immunopeptidomic landscape of ovarian carcinomas. Proc National Acad Sci 114, E9942–E9951 (2017).

43. Löffler, M. W. et al. Mapping the HLA ligandome of Colorectal Cancer Reveals an Imprint of Malignant Cell Transformation. Cancer Res 78, canres.1745.2017 (2018).

44. Schmidt, J. et al. Prediction of neo-epitope immunogenicity reveals TCR recognition determinants and provides insight into immunoediting. Cell Reports Medicine 2, 100194 (2021).

45. Wells, D. K. et al. Key Parameters of Tumor Epitope Immunogenicity Revealed Through a Consortium Approach Improve Neoantigen Prediction. Cell 183, 818–834.e13 (2020).

46. Bassani-Sternberg, M. et al. Direct identification of clinically relevant neoepitopes presented on native human melanoma tissue by mass spectrometry. Nat Commun 7, 13404 (2016).

47. Bassani-Sternberg, M. et al. Deciphering HLA-I motifs across HLA peptidomes improves neo-antigen predictions and identifies allostery regulating HLA specificity. Plos Comput Biol 13, e1005725 (2017).

48. Gonzalez-Galarza, F. F. et al. Allele frequency net database (AFND) 2020 update: gold-standard data classification, open access genotype data and new query tools. Nucleic Acids Res 48, D783–D788 (2020).

49. Shao, X. M. et al. High-Throughput Prediction of MHC Class I and II Neoantigens with MHCnuggets. Cancer Immunol Res 8, 396–408 (2020).

50. Ma, B. et al. PEAKS: powerful software for peptide de novo sequencing by tandem mass spectrometry. Rapid Commun Mass Sp 17, 2337–2342 (2003).

51. Frankish, A. et al. GENCODE 2021. Nucleic Acids Res 49, D916–D923 (2020).

52. Wu, T. D., Reeder, J., Lawrence, M., Becker, G. & Brauer, M. J. Statistical Genomics, Methods and Protocols. Methods Mol Biology Clifton N J 1418, 283–334 (2016).

53. Pau, G. & Reeder, J. HTSeqGenie: A NGS analysis pipeline. R package version 4.25.1. (2021).

54. Patro, R., Duggal, G., Love, M. I., Irizarry, R. A. & Kingsford, C. Salmon provides fast and bias-aware quantification of transcript expression. Nat Methods 14, 417–419 (2017).

55. Carbon, S. et al. The Gene Ontology resource: enriching a GOld mine. Nucleic Acids Res 49, D325–D334 (2020).

56. Pearson, H. et al. MHC class I-associated peptides derive from selective regions of the human genome. J Clin Invest 126, 4690–4701 (2016).

57. Cunningham, F. et al. Ensembl 2022. Nucleic Acids Res 50, D988–D995 (2021).

58. Tareen, A. & Kinney, J. B. Logomaker: beautiful sequence logos in Python. Bioinformatics 36, 2272–2274 (2019).

59. Wagih, O. ggseqlogo: a versatile R package for drawing sequence logos. Bioinform Oxf Engl 33, 3645–3647 (2017).

60. Paszke, A. et al. PyTorch: An Imperative Style, High-Performance Deep Learning Library. Arxiv (2019).

61. Howard, J. & Gugger, S. fastai: A Layered API for Deep Learning. Arxiv (2020) doi:10.3390/info11020108.

62. Kingma, D. P. & Ba, J. Adam: A Method for Stochastic Optimization. Arxiv (2014).

63. Ramachandran, P., Zoph, B. & Le, Q. V. Searching for Activation Functions. Arxiv (2017).

64. Nielsen, M. et al. NetMHCpan, a Method for Quantitative Predictions of Peptide Binding to Any HLA-A and -B Locus Protein of Known Sequence. Plos One 2, e796 (2007).

65. Henikoff, S. & Henikoff, J. G. Amino acid substitution matrices from protein blocks. Proc National Acad Sci 89, 10915–10919 (1992).

66. GitHub - facebookresearch/esm: Evolutionary Scale Modeling (esm): Pretrained language models for proteins. https://github.com/facebookresearch/esm.

67. Tareen, A. & Kinney, J. B. Logomaker: beautiful sequence logos in Python. Bioinformatics 36, 2272–2274 (2019).

